# DIPTAR: A synthetic biology platform for functional interrogation of protein degradation

**DOI:** 10.64898/2026.08.26.747229

**Authors:** Patrick M. Exconde, William Yoo, Madhura Kulkarni, Ashutosh B. Mahale, Benjamin E. Myers, Robert C. Patio, Christopher M. Bourne, Bohdana M. Discher, Cornelius Y. Taabazuing

## Abstract

Protein degradation regulates cellular homeostasis, yet many degradation events are difficult to study because they lack a readily selectable phenotype. Here, we develop Degradation-Induced Pyroptosis TArgeting Receptors (DIPTAR), a modular synthetic biology platform that couples protein degradation to CARD8-mediated pyroptosis. Using HIF-1α as a model substrate, we show that DIPTAR faithfully reports oxygen-dependent VHL-mediated degradation and enables pooled CRISPR screening to identify established and previously unrecognized regulators of HIF-1α stability. DIPTAR is functional across multiple cell types and can be programmed with diverse proteins, including BRD4, IκBα, and p53, to convert distinct degradation stimuli into a common pyroptotic output. DIPTAR also detects pathogen-mediated perturbations of host degradation pathways, including both inhibition and induction of degradation-dependent signaling. By converting protein degradation into a robust selectable phenotype, DIPTAR provides a scalable platform for functional genetic discovery, interrogation of degradation pathways, degrader characterization, and investigation of host-pathogen interactions.

## Introduction

Protein degradation is essential for cellular homeostasis, and its dysregulation contributes to cancer, neurodegeneration, infection, and other diseases^1^. This has driven the development of targeted protein degradation (TPD), in which heterobifunctional molecules recruit E3 ubiquitin ligases to disease-associated proteins to promote their degradation^2,3^. Despite considerable therapeutic promise, current TPD strategies rely predominantly on two E3 ligases, cereblon (CRBN) and von Hippel-Lindau (VHL), although the human genome encodes more than 600 E3 ligases^4^. Identifying additional substrate-E3 ligase relationships could therefore expand the therapeutic scope of TPD. Furthermore, protein degradation is also increasingly recognized as an important facet of host-pathogen interactions, as pathogens can exploit host degradation pathways to eliminate immune proteins and facilitate immune evasion^5-8^.

The ubiquitin-proteasome system (UPS) mediates approximately 80% of intracellular protein turnover in mammalian cells, yet many degradation events are difficult to detect, particularly when they lack an observable phenotype^9^. Existing approaches, including reporter assays, proteomics, and genetic screens, often provide indirect or low-throughput readouts, require specialized instrumentation, or lack a robust selectable phenotype for identifying regulators of protein turnover. Approaches that directly link protein degradation to a functional cellular outcome could facilitate mechanistic studies of proteostasis and accelerate the discovery of therapeutically relevant degradation pathways.

The mammalian innate immune system uses pattern-recognition receptors (PRRs) to detect invading pathogens and their activities^10^. Some PRRs trigger pyroptosis, a lytic form of programmed cell death that protects against infection^11,12^. Caspase recruitment domain-containing protein 8 (CARD8) is an intracellular PRR that undergoes post-translational autoproteolysis to generate two non-covalently associated fragments (Fig. 1A)^13^. In response to stimuli such as HIV protease activity or the dipeptidyl peptidase inhibitor Val-boroPro, the auto-proteolyzed CARD8 N-terminal fragment undergoes ubiquitin-independent proteasomal degradation, releasing the C-terminal fragment to assemble an inflammasome (Fig. 1A)^14-17^. Inflammasome formation activates caspase-1 (CASP1), which processes the pro-inflammatory cytokines IL-1β and IL-18 and cleaves gasdermin D (GSDMD) to induce pyroptosis^18^. Thus, CARD8 provides a natural mechanism for converting protein degradation into cell death.

**Fig. 1.**
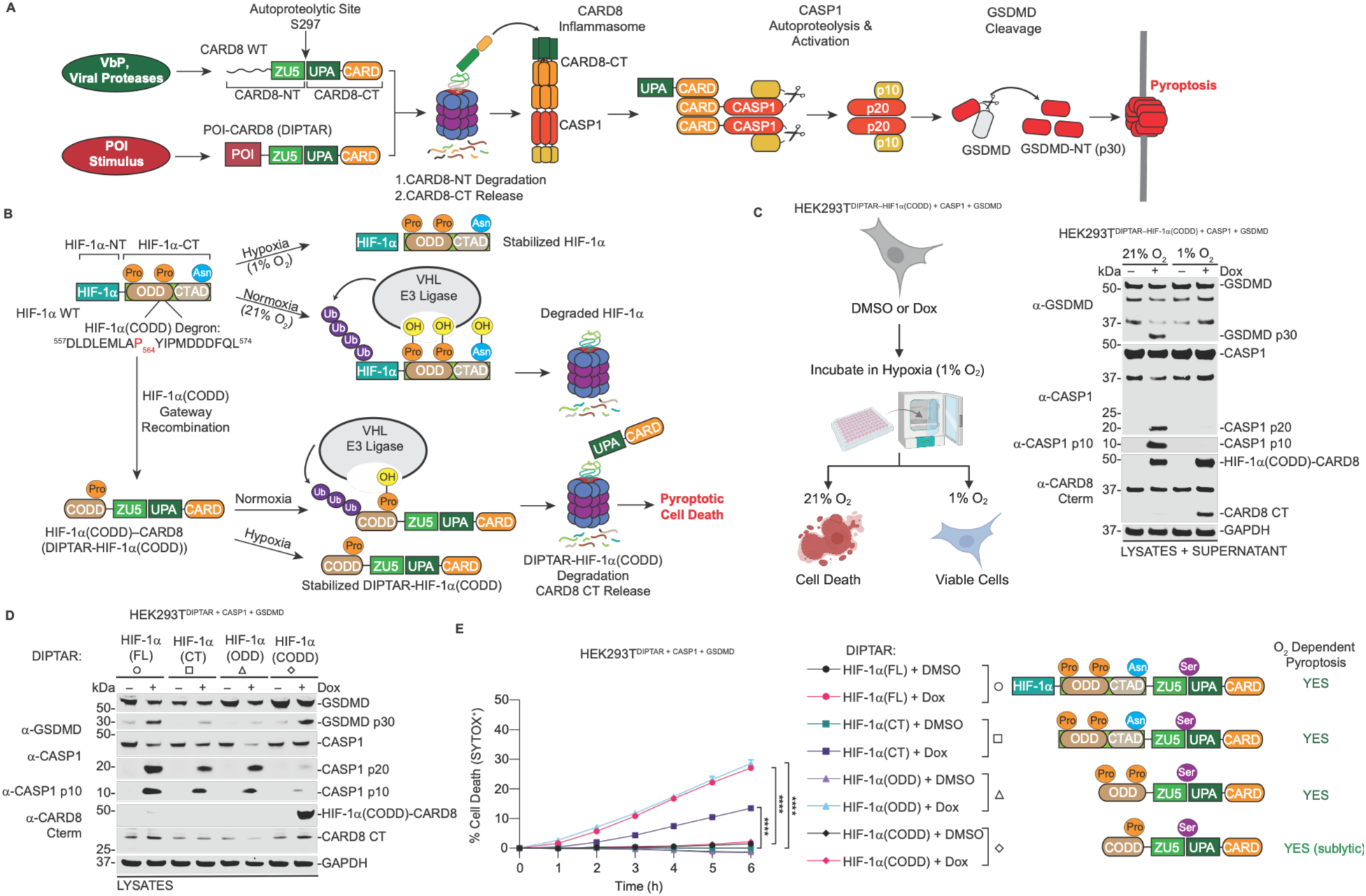
DIPTAR converts oxygen-dependent HIF-1α degradation into pyroptotic cell death. (**A**) Schematic of WT CARD8 and POI-CARD8 receptors. N-terminal degradation releases the CARD8 C-terminal fragment, promoting inflammasome formation, CASP1 activation, GSDMD cleavage, and pyroptosis. (**B**) Schematic of oxygen-dependent HIF-1α degradation and design of the HIF-1α-CARD8 receptor. (**C**) Experimental design (left) and immunoblot analysis of HEK293T cells stably expressing CASP1, GSDMD, and DIPTAR-HIF-1α(CODD). Cells were treated with DMSO or doxycycline (Dox; 1 μg/mL) and incubated under hypoxia (1% O_2_) for 12 h, then either maintained under hypoxia or reoxygenated at 21% O_2_ for 1 h before analysis. (**D,E**) HEK293T cells expressing CASP1, GSDMD, and the indicated HIF-1α-based DIPTARs were treated with Dox (1 μg/mL) under hypoxia for 12 h and then reoxygenated at 21% O_2_. (**D**) Cells were reoxygenated for 1 h before immunoblot analysis. (**E**) Cell-death kinetics following reoxygenation were measured by SYTOX Green uptake using an Incucyte S3. Data are mean ± SEM of three independent biological experiments. ****P < 0.0001, ***P < 0.001, **P < 0.01, and *P < 0.05 by two-way ANOVA with Tukey’s multiple-comparisons test.

We reasoned that fusing proteins of interest (POIs) to CARD8 N-terminus could exploit this mechanism to create synthetic receptors that convert POI degradation into a selectable phenotype (Fig. 1A). Here, we develop <u>D</u>egradation-Induced <u>P</u>yroptosis <u>TA</u>rgeting <u>R</u>eceptors (DIPTAR), a modular synthetic biology platform that couples degradation of a POI to CARD8-mediated pyroptosis. Using Hypoxia-inducible factor 1α (HIF-1α) as a model POI, we show that DIPTAR reports on oxygen-dependent degradation and enables functional genetic discovery of regulators of protein stability. We further demonstrate that the DIPTAR platform is adaptable across diverse POIs and cellular contexts and can also report pathogen-mediated modulation of host protein degradation. Together, these findings establish DIPTAR as a scalable platform for functional interrogation of protein degradation pathways.

## Results

### DIPTAR converts oxygen-dependent HIF-1α degradation into pyroptotic cell death

Hypoxia-inducible factor 1α (HIF-1α) is a transcription factor that is regulated by oxygen-dependent degradation. Under normoxia (21% O_2_), HIF-1α is hydroxylated at prolines 402 and 564, promoting recognition by the von Hippel-Lindau (VHL) E3 ligase complex and subsequent proteasomal degradation (Fig. 1B)^19^. Under hypoxia (1% O_2_), hydroxylation is inhibited, resulting in HIF-1α stabilization and transcriptional activation^20,21^. To determine whether CARD8 could be reprogrammed to sense oxygen-dependent protein degradation, we fused the C-terminal oxygen-dependent degradation domain (CODD) of HIF-1α, which contains proline 564, to the CARD8 N-terminus. To facilitate modular receptor engineering, we generated inducible and constitutive Gateway-compatible vectors that fuse the C-terminus of a POI to the N-terminus of CARD8 beginning at residue 151 (Fig. S1A). We termed this platform <u>D</u>egradation-Induced <u>P</u>yroptosis <u>TA</u>rgeting <u>R</u>eceptors (DIPTAR) and refer to individual receptors as DIPTAR-POI. Thus, the HIF-1α(CODD)-CARD8 receptor is designated DIPTAR-HIF-1α(CODD). We reasoned that this receptor would remain stable under hypoxia but undergo degradation and induce pyroptosis upon reoxygenation (Fig. 1B).

To test this, we generated HEK293T cells stably expressing CASP1, GSDMD and doxycycline inducible DIPTAR-HIF-1α(CODD). Receptor expression was induced under hypoxia for 12 h, followed by reoxygenation for 1 h. Because HIF-1α has a short half-life following reoxygenation (∼3–5 min)^22^, hypoxic samples were rapidly processed to minimize unintended oxygen exposure. Reoxygenation resulted in DIPTAR-HIF-1α(CODD) degradation, CASP1 activation, and GSDMD cleavage, demonstrating that DIPTAR can couple oxygen-dependent protein degradation to pyroptotic signaling (Fig. 1C).

We next examined whether DIPTAR could accommodate POIs of varying size. We generated receptors containing full-length HIF-1α (FL), the C-terminal region (CT), or the oxygen-dependent degradation domain (ODD), all of which retain both VHL-recognition prolines. Each receptor induced CASP1 activation and GSDMD cleavage (Fig. 1D). The FL and ODD receptors induced robust lytic cell death as assessed by SYTOX Green uptake, whereas the CT receptor, which exhibited lower expression (Fig. S1B), produced a weaker response (Fig. 1E). In contrast, the CODD receptor induced GSDMD cleavage without detectable lytic cell death, suggesting that its degradation generates sufficient signaling for GSDMD processing but does not reach the threshold required for overt pyroptosis (Fig. 1D,E). Together, these findings demonstrate that DIPTAR can accommodate POIs ranging from approximately 2 to 100 kDa and convert their degradation into pyroptotic signaling.

### DIPTAR-HIF-1α-induced pyroptosis requires the VHL degron, proteasomal degradation, and caspase-1 activation

To define the mechanism of HIF-1α DIPTAR-induced pyroptosis, we focused on the DIPTAR-HIF-1α(ODD) receptor, which was sufficient to induce robust cell death. We generated an autoproteolysis-deficient CARD8 mutant (S297A), a degron-deficient mutant in which the VHL-recognition prolines were mutated to alanine (P402A/P564A), and control receptors containing enhanced green fluorescent protein (eGFP) or the HIF-1α C-terminal transactivation domain (CTAD), which undergoes oxygen-dependent hydroxylation but is not targeted for VHL-mediated degradation (Fig. 2A). Only DIPTAR-HIF-1α(ODD) induced CASP1 activation, GSDMD cleavage, and SYTOX Green uptake following reoxygenation, demonstrating that pyroptosis requires both a functional VHL-recognition degron and CARD8 autoproteolysis (Fig. 2A,B). Whereas DIPTAR-HIF-1α(ODD) expression was largely undetectable by immunoblotting, the degron-deficient, eGFP, and CTAD receptors accumulated to higher levels, consistent with impaired degradation.

**Fig. 2.**
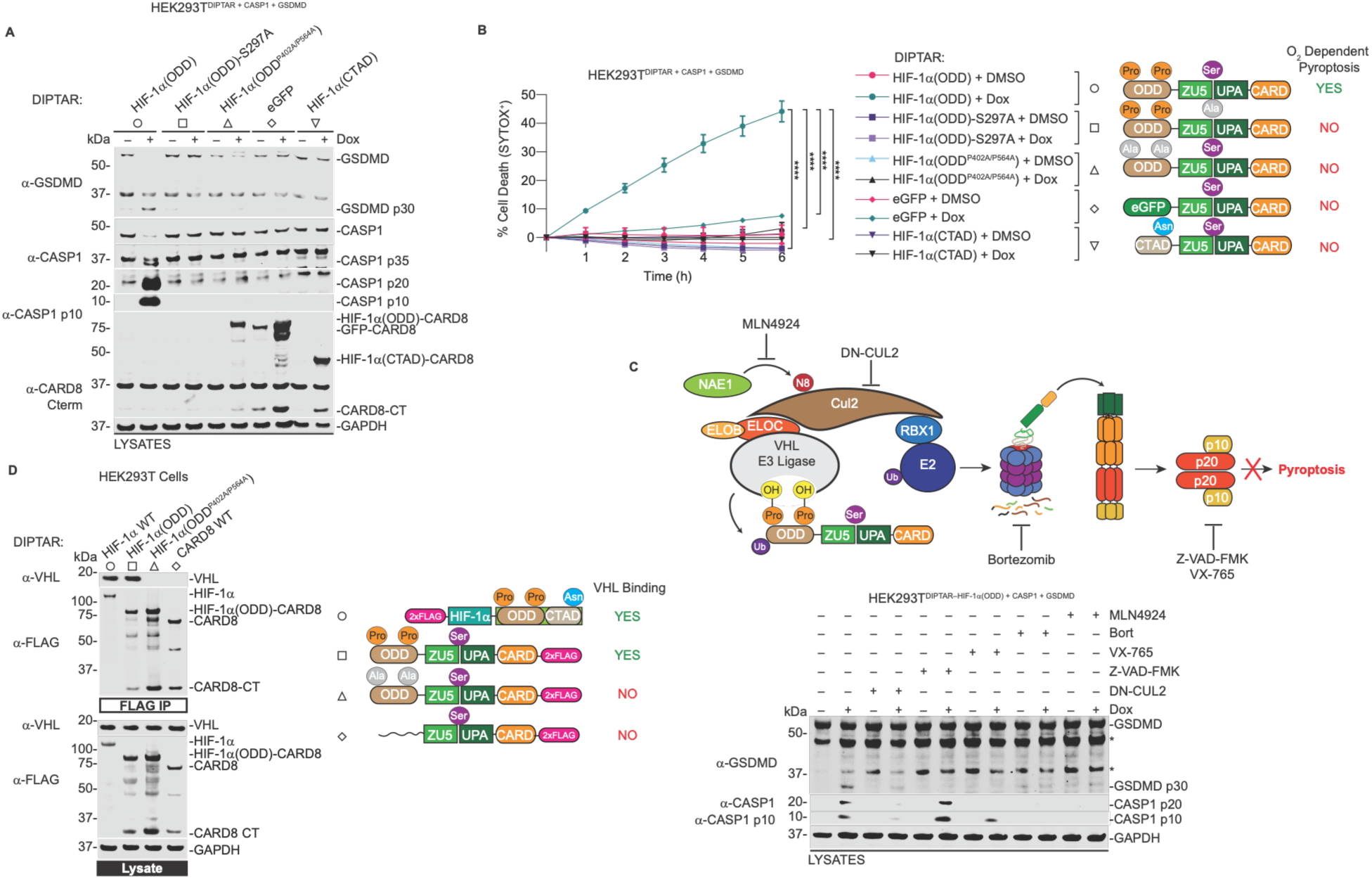
DIPTAR-HIF-1α-induced pyroptosis requires the VHL degron, proteasomal degradation, and caspase-1 activation. (**A,B**) HEK293T cells stably expressing CASP1, GSDMD, and the indicated DIPTAR-HIF-1α(ODD) variants or controls were treated with Dox (1 μg/mL) under hypoxia (1% O_2_) for 12 h and then reoxygenated at 21% O_2_. (**A**) Cells were reoxygenated for 1 h before immunoblot analysis. (**B**) Cell-death kinetics following reoxygenation were measured by SYTOX Green uptake using an Incucyte S3. (**C**) HEK293T cells stably expressing CASP1, GSDMD, and DIPTAR-HIF-1α(ODD) were transfected with a plasmid encoding dominant-negative CUL2 (DN-CUL2; 100 ng) or pretreated for 30 min with Z-VAD-FMK (40 μM), VX-765 (40 μM), MLN4924 (1 μM), or bortezomib (1 μM). Cells were incubated under hypoxia for 6 h and then reoxygenated for 1 h before immunoblot analysis. The mechanisms of action of the indicated inhibitors are shown above. (**D**) HEK293T cells were transfected with the indicated FLAG-tagged constructs for 48 h, followed by FLAG immunoprecipitation and immunoblot analysis of endogenous VHL association. Data are mean ± SEM of three independent biological experiments. ****P < 0.0001, ***P < 0.001, **P < 0.01, and *P < 0.05 by two-way ANOVA with Tukey’s multiple-comparisons test.

We next inhibited components of the canonical HIF-1α degradation pathway. VHL functions within a Cullin-RING E3 ligase complex containing CUL2, ELOB, ELOC, and RBX1 (Fig. 2C)^23^. Expression of a dominant-negative CUL2 construct, inhibition of cullin neddylation with MLN4924^24^, or proteasome inhibition with bortezomib^25^ abolished CASP1 activation and GSDMD cleavage. Similarly, the caspase inhibitors Z-VAD-FMK and VX-765^26^ blocked GSDMD cleavage (Fig. 2C). These findings demonstrate that DIPTAR-HIF-1α-induced pyroptosis requires intact Cullin-RING ligase activity, proteasomal degradation, and CASP1 activity.

To determine whether DIPTAR-HIF-1α engages VHL, we performed FLAG co-immunoprecipitation experiments using wild-type (WT) HIF-1α, DIPTAR-HIF-1α(ODD), DIPTAR-HIF-1α(ODD^P402A/P564A^), and wild-type CARD8. WT HIF-1α and DIPTAR-HIF-1α(ODD) efficiently co-immunoprecipitated endogenous VHL, whereas the degron-deficient receptor and WT CARD8 did not (Fig. 2D). Consistent with these results, all HIF-1α-based DIPTARs containing one or both VHL-recognition prolines, including FL, CT, ODD, and CODD, co-immunoprecipitated endogenous VHL (Fig. S2A). Collectively, these data demonstrate that DIPTAR-HIF-1α preserves canonical VHL-dependent recognition and converts oxygen-dependent proteasomal degradation into CASP1-dependent pyroptotic signaling.

### DIPTAR function is preserved across multiple cell types

Because E3 ligase expression and activity vary across tissues and cell types, we next examined whether DIPTAR functions beyond HEK293T cells. We generated stable DIPTAR-HIF-1α(ODD) cell lines in THP-1 monocytes and HCC1806 epithelial cells, both of which endogenously express CASP1 and GSDMD. Upon reoxygenation, differentiated THP-1 macrophages exhibited robust SYTOX Green uptake, CASP1 activation, and cleavage of GSDMD and IL-1β (Fig. 3A,B). These responses were abolished by Z-VAD-FMK or VX-765, supporting CASP1-dependent pyroptosis.

**Fig. 3.**
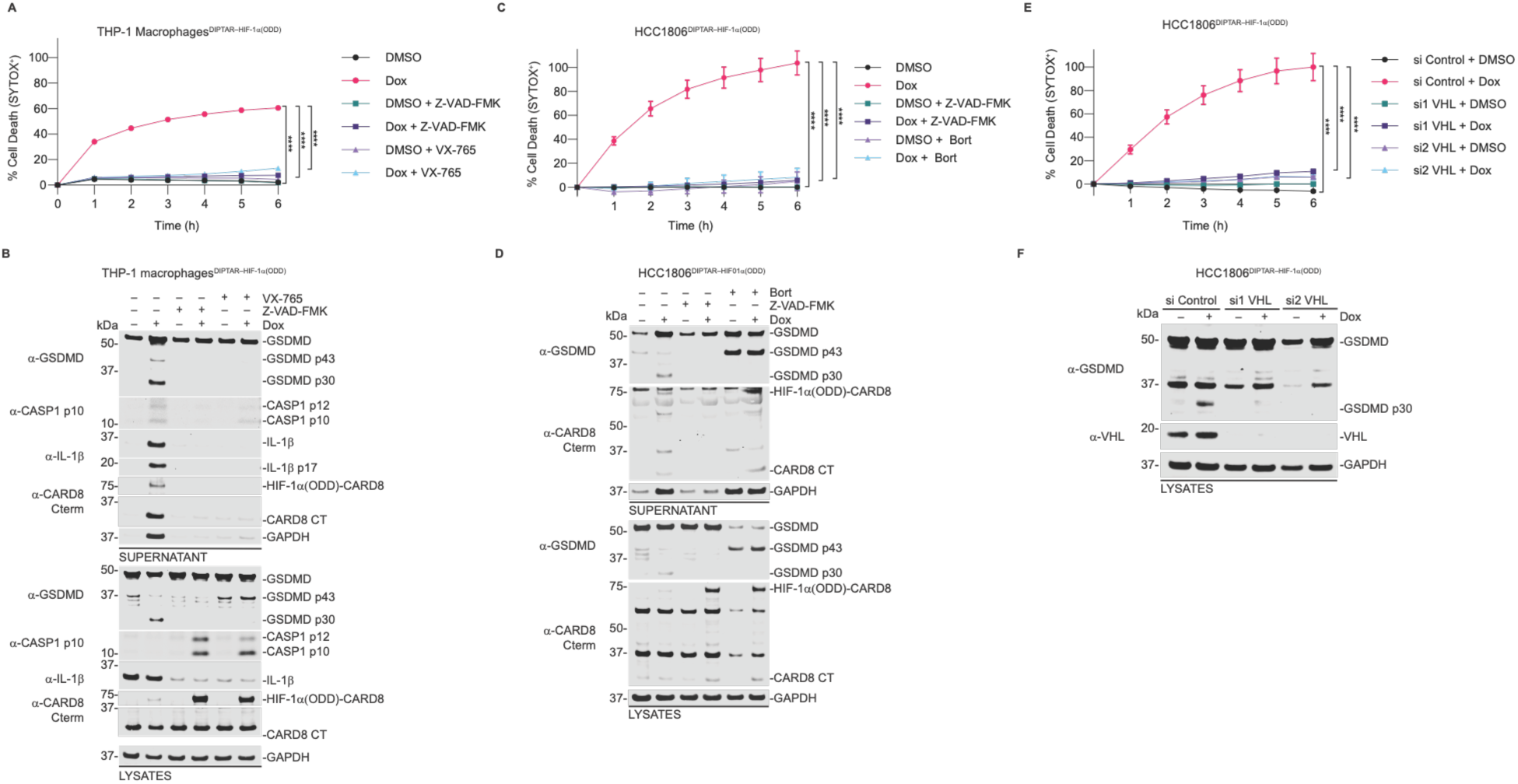
DIPTAR function is preserved across multiple cell types. (**A,B**) PMA-differentiated THP-1 macrophages stably expressing DIPTAR-HIF-1α(ODD) were pretreated with Z-VAD-FMK or VX-765 (40 μM) for 30 min and induced with Dox (1 μg/mL) under hypoxia for 12 h before reoxygenation. Cell-death kinetics were measured by SYTOX Green uptake (**A**), and terminal samples were analyzed by immunoblotting (**B**). (**C,D**) HCC1806 cells stably expressing DIPTAR-HIF-1α(ODD) were treated with Z-VAD-FMK (40 μM) or bortezomib (50 nM) and Dox (1 μg/mL) under hypoxia for 12 h before reoxygenation. Cell-death kinetics were measured by SYTOX Green uptake (**C**), and terminal samples were analyzed by immunoblotting (**D**). (**E,F**) HCC1806 cells stably expressing DIPTAR-HIF-1α(ODD) were transfected with two independent VHL-targeting siRNAs for 24 h under normoxia, induced with Dox (1 μg/mL) under hypoxia for 12 h, and then reoxygenated. Cell-death kinetics were measured by SYTOX Green uptake (**E**), and terminal samples were analyzed by immunoblotting (**F**). Data are mean ± SEM of three independent biological experiments. ****P < 0.0001, ***P < 0.001, **P < 0.01, and *P < 0.05 by two-way ANOVA with Tukey’s multiple-comparisons test.

Similarly, reoxygenation induced SYTOX Green uptake and GSDMD cleavage in HCC1806 cells expressing DIPTAR-HIF-1α(ODD), both of which were suppressed by bortezomib or Z-VAD-FMK (Fig. 3C,D). To confirm dependence on the canonical oxygen-sensing pathway, we depleted VHL using siRNA. VHL depletion reduced both SYTOX Green uptake and GSDMD cleavage relative to the non-targeting control (Fig. 3E,F). Together, these findings demonstrate that DIPTAR can couple protein degradation to pyroptosis across distinct immune and epithelial cell types using endogenous pyroptotic machinery.

### DIPTAR couples protein degradation to functional genetics

Because DIPTAR converts protein degradation into a selectable cell-death phenotype, we reasoned that it could be adapted for pooled CRISPR screening to identify regulators of protein turnover. To enable this approach, we engineered a DIPTAR vector containing a U6 promoter-sgRNA expression cassette downstream of CARD8, enabling co-expression of the receptor and sgRNA within the same cell (Fig. S3A). HCC1806 cells stably expressing Cas9 (Fig. S3B) were transduced with DIPTAR-HIF-1α(ODD) carrying the focused Bison sgRNA library, which targets 713 E1, E2, and E3 ligases, deubiquitinating enzymes (DUBs), and control genes with 2,852 sgRNAs total^27^. Cells were transduced at low multiplicity of infection to maintain single-guide representation and library complexity (Fig. S3C). Following selection, cells were maintained under hypoxia to induce receptor expression and subsequently reoxygenated. Because disruption of genes required for HIF-1α degradation should prevent DIPTAR-induced pyroptosis, sgRNAs targeting positive regulators of degradation are expected to become enriched among surviving cells (Fig. 4A and Fig. S3C).

**Fig. 4.**
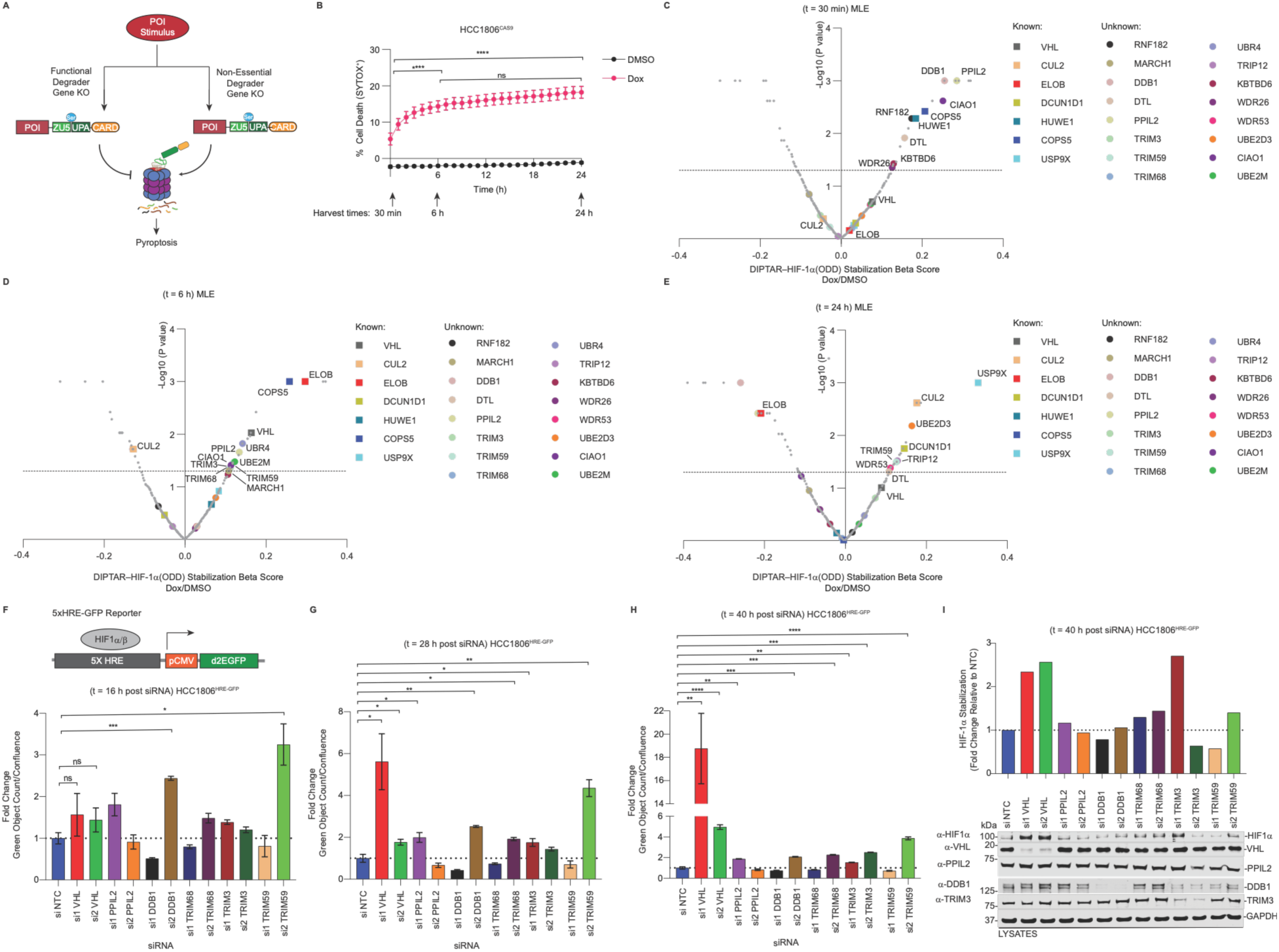
DIPTAR couples protein degradation to functional genetics. (**A**) Schematic of the DIPTAR-based CRISPR screening strategy. Knockout of genes required for POI degradation prevents DIPTAR activation and cell death, enriching the corresponding sgRNAs in surviving cells. (**B**) HCC1806-Cas9 cells were transduced with a DIPTAR vector encoding HIF-1α(ODD)-CARD8 and the focused Bison sgRNA library. Cell-death kinetics were monitored, and cells were collected at 30 min, 6 h, and 24 h for next-generation sequencing. (**C–E**) Volcano plots showing genes enriched in surviving cells at 30 min (**C**), 6 h (**D**), and 24 h (**E**). (**F–H**) HCC1806 cells stably expressing an HRE-driven GFP reporter were transfected with siRNAs targeting the indicated screen hits under normoxic conditions. At the indicated times after siRNA transfection, GFP-positive objects were quantified and normalized to cell confluence using an Incucyte S3. GFP signal is shown as fold change relative to the non-targeting control (NTC) siRNA. (**I**) Immunoblot analysis of HIF-1α in siRNA-treated cells at the terminal time point. HIF-1α band intensity was quantified by densitometry, normalized to GAPDH, and expressed as fold change relative to NTC-treated cells. Data in **F–H** are mean ± SEM of three independent biological experiments. ****P < 0.0001, ***P < 0.001, **P < 0.01, and *P < 0.05 by a two tailed Student’s *t*-test.

To capture temporal differences in degradation pathways, we first characterized cell-death kinetics and collected cells at 30 min, 6 h, and 24 h following reoxygenation (Fig. 4B). sgRNA enrichment was analyzed using MAGeCK maximum-likelihood estimation (MLE) and robust rank aggregation (RRA) (Fig. 4C-E and Fig. S3D,E), which yielded similar results^28,29^. Distinct patterns of enrichment emerged over time. Early hits included *HUWE1* and *COPS5*, which have established roles in HIF-1α regulation outside the canonical oxygen-dependent VHL pathway (Fig. 4C)^30,31^. In contrast, later time points enriched components of the canonical oxygen-dependent degradation machinery, including *VHL*, *ELOB*, *CUL2*, and *DCUN1D1* (Fig. 4D,E)^23,32,33^, supporting the specificity and temporal resolution of the screen. Other candidates displayed delayed enrichment, such as *USP9X*, which stabilizes the E3 ligase SMURF1, a reported regulator of VHL stability^34^.

The screen also identified factors with plausible but incompletely defined connections to HIF-1α turnover. These included *CIAO1*, a regulator of iron homeostasis that could influence the iron-dependent hydroxylases controlling HIF-1α stability^35^; *UBE2M*, the NEDD8-conjugating E2 that regulates cullin neddylation^36^; and *UBE2D3*, a member of an E2 family implicated in VHL-mediated HIF-α ubiquitination, although a specific role for UBE2D3 in HIF-1α turnover has not been established^37^. In addition, the screen identified candidate regulators not previously linked to HIF-1α degradation, including *RNF182, PPIL2, DDB1, TRIM3, TRIM59,* and *TRIM68* (Fig. 4C-E). To validate candidate regulators independently of DIPTAR, we generated HCC1806 cells expressing an HRE-GFP reporter, in which HIF-1α stabilization increases GFP expression (Fig. 4F). We prioritized candidates without established roles in HIF-1α regulation and depleted each using two independent siRNAs under normoxic conditions. DDB1 and TRIM59 depletion increased GFP signal approximately threefold by 16 h, when VHL depletion produced little detectable increase (Fig. 4G,H). However, the effects of the candidate regulators remained modest over time, whereas VHL depletion progressively increased GFP signal to approximately 6-fold at 28 h and 20-fold at 40 h relative to the non-targeting control. Immunoblotting at 40 h confirmed increased HIF-1α protein following depletion of VHL, TRIM3, TRIM59, and TRIM68 (Fig. 4I). Together, these findings demonstrate that DIPTAR can be coupled to pooled functional genomics to identify regulators of protein stability. The magnitude and kinetics of the validation phenotypes further support VHL as the dominant regulator of HIF-1α degradation under normoxic conditions, while revealing additional factors that may contribute to basal HIF-1α turnover or modulate the efficiency of its degradation.

### DIPTAR enables modular engineering of receptors that couple distinct protein degradation stimuli to pyroptosis

To evaluate the modularity of DIPTAR, we generated receptors incorporating BRD4, IκBα, or p53, which undergo degradation through distinct regulatory mechanisms, and expressed them in HCC1806 cells (Fig. 5A). BRD4 is a transcriptional co-regulator that controls oncogenic programs, including MYC expression^38^, and can be targeted for degradation by the PROTAC dBET1 through recruitment of the CRBN E3 ligase complex^39^. Treatment of DIPTAR-BRD4-expressing cells with dBET1 induced robust SYTOX Green uptake and GSDMD cleavage (Fig. 5B,C). These responses were suppressed by the CRBN inhibitor EM12-SO_2_F^40^, MLN4924, bortezomib, and Z-VAD-FMK, demonstrating that DIPTAR-BRD4 couples CRBN-mediated degradation to pyroptotic signaling.

**Fig. 5.**
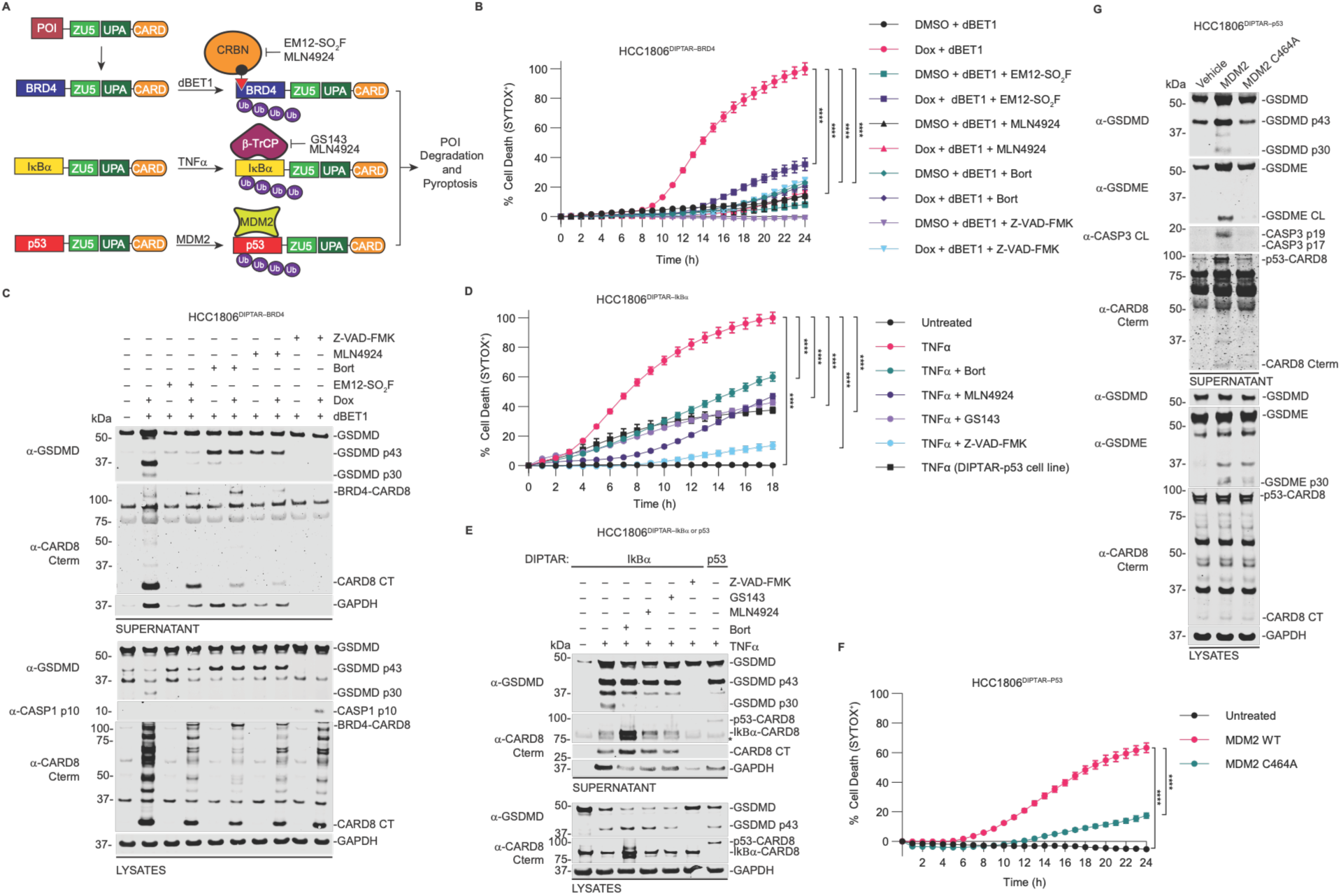
DIPTAR enables modular engineering of receptors that couple distinct protein degradation stimuli to pyroptosis. (**A**) Schematic of the indicated receptors generated using the DIPTAR platform. (**B,C**) HCC1806 cells stably expressing Dox-inducible DIPTAR-BRD4 were pretreated for 30 min with EM12-SO_2_F (1 μM), bortezomib (1 μM), MLN4924 (1 μM), or Z-VAD-FMK (40 μM), followed by dBET1 (1 μM) for 30 min before Dox induction (1 μg/mL). Cell-death kinetics were measured by SYTOX Green uptake (**B**), and terminal samples were analyzed by immunoblotting (**C**). (**D,E**) HCC1806 cells stably expressing DIPTAR-IκBα or DIPTAR-p53 were pretreated for 30 min with bortezomib (100 nM), MLN4924 (1 μM), GS143 (1 μM), or Z-VAD-FMK (40 μM), followed by TNFα (200 ng/mL) to activate NF-κB signaling. Cell-death kinetics were measured by SYTOX Green uptake (**D**), and terminal samples were analyzed by immunoblotting (**E**). (**F,G**) HCC1806 cells stably expressing DIPTAR-p53 were transfected with plasmids (50 ng) encoding WT MDM2 or catalytically inactive MDM2 (C464A). Cell-death kinetics were measured by SYTOX Green uptake (**F**), and terminal samples were analyzed by immunoblotting (**G**). Data are mean ± SEM of three independent biological experiments. ****P < 0.0001, ***P < 0.001, **P < 0.01, and *P < 0.05 by two-way ANOVA with Tukey’s multiple-comparisons test.

We next examined IκBα, a negative regulator of NF-κB signaling that undergoes rapid β-TrCP-dependent proteasomal degradation following pathway activation, permitting NF-κB nuclear translocation and transcriptional activity^41^. TNFα induced robust SYTOX Green uptake and GSDMD cleavage in cells expressing DIPTAR-IκBα but not the DIPTAR-p53 control receptor (Fig. 5D,E). Both responses were attenuated by the β-TrCP inhibitor GS-143^42^, MLN4924, bortezomib, and Z-VAD-FMK. Residual cell death persisted with several inhibitors, likely reflecting TNFα-induced apoptotic signaling through CASP8^43^. Consistent with this interpretation, CASP3-mediated GSDMD cleavage into the inactive p43 fragment^44^ was retained, but the active p30 fragment was absent under these conditions, indicating that the residual cell death was not GSDMD-dependent pyroptosis.

Finally, we assessed whether DIPTAR could report on p53 degradation. HCC1806 cells expressing DIPTAR-p53 were co-transfected with either WT MDM2, the principal E3 ligase responsible for p53 turnover^45^, or the catalytically inactive MDM2 (C464A) mutant. WT MDM2 induced robust SYTOX Green uptake and GSDMD cleavage, whereas MDM2 C464A did not (Fig. 5F,G). We also observed CASP3 activation and GSDME cleavage in cells expressing WT MDM2, suggesting engagement of additional cell-death pathways. Together, these findings demonstrate that DIPTAR can be programmed with diverse POIs to selectively couple their cognate degradation stimuli to a common pyroptotic output.

### DIPTAR enables functional detection of pathogen effector activity

Many pathogens manipulate host proteostasis to evade immune detection and promote infection by utilizing effectors that target host proteins for degradation^5-8,46,47^. Because these events can be difficult to detect and functionally characterize, we investigated whether DIPTAR could report pathogen-mediated modulation of host protein degradation.

The intracellular bacterial pathogen *Coxiella burnetii* injects effector proteins that suppress host immune signaling and facilitate intracellular replication. One such effector, CBU1314, has been proposed to inhibit NF-κB signaling by suppression of IKK activation (Fig. 6A)^48^. To test whether DIPTAR could detect this activity, HCC1806 cells expressing DIPTAR-IκBα were stimulated with TNFα in the presence or absence of CBU1314. TNFα induced robust propidium iodide uptake and GSDMD and GSDME cleavage, whereas CBU1314 markedly attenuated these responses (Fig. 6B,C), consistent with suppression of NF-κB signaling upstream of IκBα degradation.

**Fig. 6.**
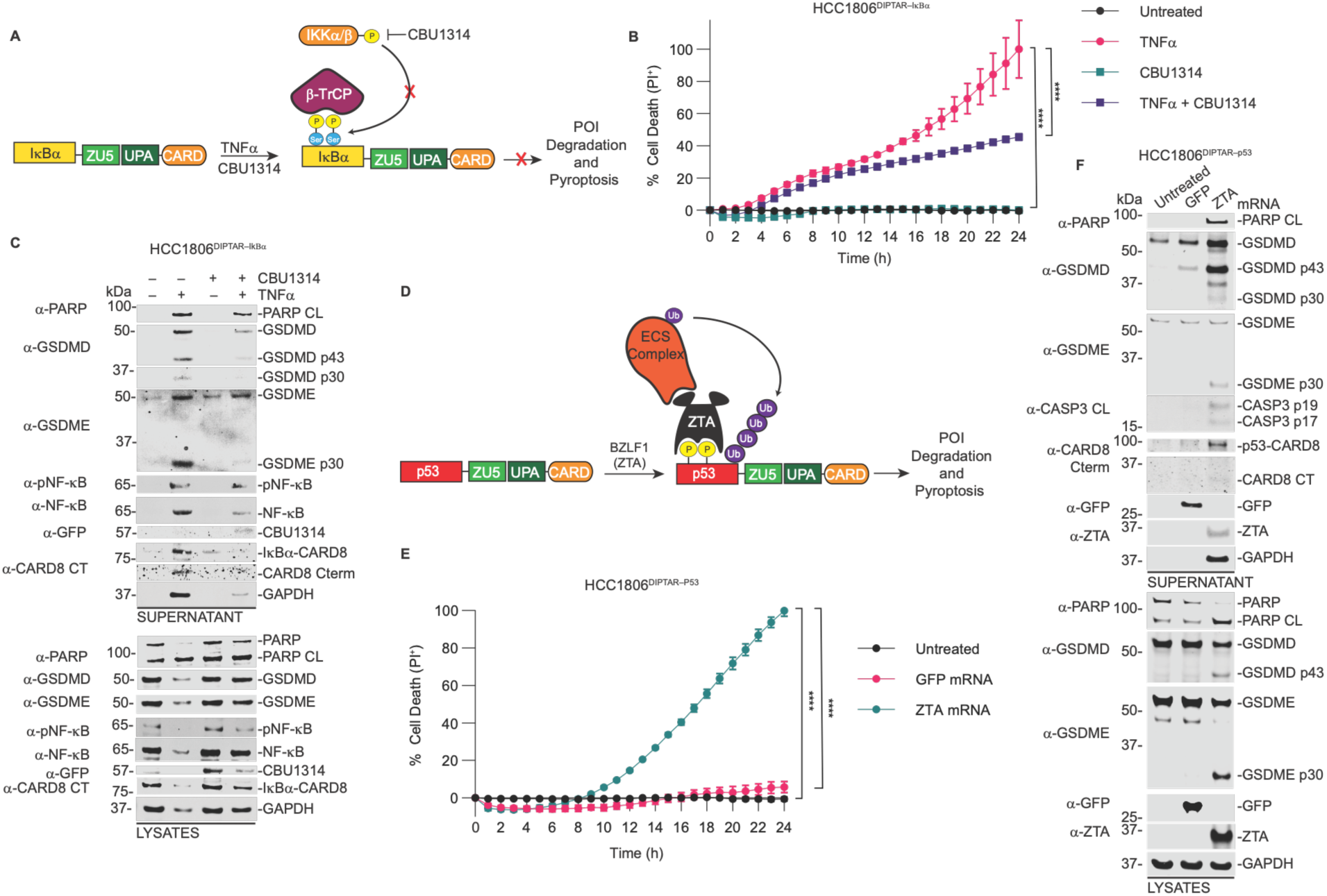
DIPTAR enables functional detection of pathogen effector activity. (**A**) Schematic of the DIPTAR-IκBα receptor and its activation through IκBα degradation. (**B,C**) HCC1806 cells stably expressing DIPTAR-IκBα were transfected with a plasmid (200 ng) encoding the *Coxiella burnetii* effector CBU1314 and treated with TNFα (100 ng/mL) 30 min later. Cell-death kinetics were measured by PI uptake (**B**), and terminal samples were analyzed by immunoblotting (**C**). (**D**) Schematic of the DIPTAR-p53 receptor and its activation through p53 degradation. (**E,F**) HCC1806 cells stably expressing DIPTAR-p53 were transfected with mRNA encoding GFP or the Epstein-Barr virus effector ZTA. Cell-death kinetics were measured by PI uptake (**E**), and terminal samples were analyzed by immunoblotting (**F**). Data are mean ± SEM of three independent biological experiments. ****P < 0.0001, ***P < 0.001, **P < 0.01, and *P < 0.05 by two-way ANOVA with Tukey’s multiple-comparisons test.

We next examined Epstein-Barr virus (EBV), which alters host p53 regulation during viral reactivation^49^. The EBV protein BZLF1 (ZTA) has been implicated in recruiting the host ECS (Elongin B/C–Cullin2/5–SOCS-box) degradation machinery to regulate p53 stability (Fig. 6D)^49^. HCC1806 cells expressing DIPTAR-p53 were transfected with in vitro-transcribed GFP or ZTA mRNA. Whereas GFP had little effect, ZTA induced robust propidium iodide uptake and GSDMD and GSDME cleavage, likely due to CASP1-mediated activation of CASP3 which subsequently cleaves GSDME (Fig. 6E,F)^50,51^. These findings are consistent with ZTA-mediated destabilization of p53 and activation of DIPTAR-p53. Altogether, these findings establish DIPTAR as a functional reporter of pathogen effector activity and demonstrate its ability to convert pathogen-mediated perturbations of host protein stability into measurable cellular phenotypes. DIPTAR therefore provides a framework for identifying otherwise cryptic pathogen-induced degradation events in living cells.

## Discussion

Protein degradation governs nearly every aspect of cellular physiology, yet degradation events are often difficult to detect unless they produce an observable phenotype. Existing approaches typically rely on biochemical measurements, fluorescent reporters, flow cytometry, or proteomic analyses that require specialized instrumentation, can be difficult to scale, and may fail to capture transient or low-abundance degradation events in living cells. Here, we developed DIPTAR, a modular synthetic biology platform that converts protein degradation into a robust and selectable cell-death phenotype. Unlike reporter-based systems that infer degradation from changes in fluorescence or protein abundance, DIPTAR couples degradation directly to cell fate. This architecture enables enrichment-based screening without fluorescence-activated cell sorting, high-content imaging, or deep proteomic profiling. Moreover, amplification through inflammasome signaling may facilitate detection of degradation events that are difficult to resolve using conventional approaches.

DIPTAR is particularly well suited for functional genetic discovery. Pooled CRISPR screening recovered core components of the canonical VHL pathway while also identifying candidate regulators not previously linked to HIF-1α stability. Independent validation revealed substantially greater HIF-1α stabilization following VHL depletion than depletion of the additional candidates, consistent with VHL serving as the dominant regulator of HIF-1α degradation under normoxic conditions while additional factors may contribute to or modulate basal turnover. Importantly, because DIPTAR reports productive degradation rather than protein binding or ubiquitination alone, it can identify both core degradation machinery and cellular pathways that influence protein turnover. This approach may therefore facilitate discovery of substrate-specific E3 ligases and regulatory pathways that could be leveraged for targeted protein degradation (TPD). Notably, we observed no apparent size limitation among the POIs tested, and DIPTAR receptors retained the biological functions and degradation responses of their cognate proteins.

Recently, the Death FUSion Escaper (DEFUSE) platform was reported, which couples protein degradation to cell survival^52^. DEFUSE fuses a POI to FKBP12 and caspase-9 such that POI degradation prevents chemically induced apoptosis and permits cell survival^52^. Although both platforms convert degradation into cell fate, DIPTAR reports degradation through CARD8-dependent pyroptosis rather than protection from inducible apoptosis. DEFUSE is limited for POIs that dimerize, aggregate, or are intrinsically cytotoxic, as these properties can induce cell death independently of degradation. DIPTAR activation, by contrast, can be distinguished from nonspecific toxicity through pathway-specific biomarkers, including CASP1 activation, GSDMD cleavage, and IL-1β processing. These features may broaden the range of proteins amenable to degradation-coupled cell-death screening while providing orthogonal validation of receptor activity.

DIPTAR also offers several potential applications for degrader discovery and characterization. Individual receptors could be used to identify molecular glues, PROTACs, or other compounds that induce degradation of a specific target. Conversely, pooled libraries of DIPTAR receptors could enable dropout screens for unintended degradation events, providing a strategy to profile degrader selectivity across multiple potential substrates. DIPTAR-based chemical and genetic screens could further identify determinants of degrader sensitivity, resistance, and substrate specificity, addressing important challenges in the development of next-generation TPD therapeutics.

The modularity of DIPTAR extends beyond degrader screening. Receptors incorporating BRD4, IκBα, and p53 responded to their respective degradation stimuli, demonstrating that distinct degradation pathways can be converted into a common pyroptotic output. The DIPTAR-IκBα receptor further illustrates how degradation can serve as a proxy for pathway activity, converting NF-κB activation into an easily detectable cell-death phenotype. Similar approaches could be applied to other signaling pathways in which regulated protein turnover serves as a key molecular switch.

DIPTAR may be particularly useful for investigating host-pathogen interactions. Pathogens frequently manipulate host ubiquitin signaling and protein stability to evade immunity, yet many of these activities produce subtle phenotypes that are difficult to detect systematically. Our pathogen studies demonstrate that DIPTAR can report such perturbations in two directions. The *Coxiella burnetii* effector CBU1314 suppressed DIPTAR-IκBα activation by inhibiting signaling upstream of IκBα degradation, whereas the Epstein-Barr virus effector ZTA induced DIPTAR-p53 signaling in a manner consistent with p53 destabilization. Thus, DIPTAR can detect both pathogen-mediated inhibition and induction of host protein degradation pathways. Coupling libraries of pathogen effectors with panels of DIPTAR receptors could enable systematic mapping of host proteins and pathways manipulated by diverse pathogens, providing a framework for defining mechanisms of immune evasion and host defense.

Several limitations of the current DIPTAR architecture remain. Because DIPTAR relies on fusion of a POI to CARD8, subcellular localization may influence receptor performance. The proteins examined here were predominantly cytosolic, and whether membrane proteins or proteins localized to other cellular compartments can be incorporated into functional DIPTAR receptors remains unknown. Engineering compartment-specific DIPTAR architectures may extend the platform to proteins in otherwise inaccessible cellular environments. CARD8 fusion could also alter folding, trafficking, or protein interactions for some substrates. Finally, because DIPTAR produces a threshold-dependent cell-death response, the magnitude of receptor activation may not scale linearly with degradation kinetics. DIPTAR is therefore best suited for functional detection and selection rather than precise quantification of protein turnover.

In summary, DIPTAR establishes a modular and scalable strategy for converting protein degradation into a selectable cellular phenotype. By enabling functional genetic discovery, interrogation of diverse degradation pathways, characterization of degrader activity, and detection of pathogen-mediated perturbations, DIPTAR provides a versatile framework for studying protein turnover in living cells. Integration with CRISPR perturbation technologies, POI and ORFeome libraries, and next-generation degrader modalities could enable functional mapping of degradation pathways at scale, accelerating efforts to define the human degradome and identify new opportunities for therapeutic intervention.

## Methods

### DIPTAR vector construction

DIPTAR vectors were generated using the pCW57.1 and pLEX307 backbones. pCW57.1 was digested with AgeI-HF, and pLEX307 was digested with EcoRV-HF and SpeI-HF. CARD8 beginning at residue E151 was cloned into each backbone using NEBuilder HiFi DNA Assembly (NEB). For immunoprecipitation experiments, a 2xFLAG tag was incorporated into the primers at the C terminus of CARD8 during assembly. The DIPTAR CRISPR screening vector was generated from the pCW57.1 DIPTAR backbone by inserting a U6 promoter-sgRNA scaffold cassette derived from pLenti-Puro using NEBuilder HiFi DNA Assembly. MluI and AvrII restriction sites were incorporated to enable sgRNA library cloning. All plasmids were verified by DNA sequencing (Azenta).

### Cloning of DIPTAR constructs

Genes of interest were cloned into pDONR221 entry vectors and transferred into DIPTAR destination vectors using Gateway cloning (Thermo Fisher Scientific) according to the manufacturer’s instructions. The resulting DIPTAR proteins contain a Gateway-derived linker (DPAFLYKVVTG or DPAFLYKVVDI) between the POI and CARD8. Point mutations were introduced using the QuikChange II Site-Directed Mutagenesis Kit (Agilent) or Q5 Site-Directed Mutagenesis Kit (NEB) according to the manufacturers’ instructions. Primers used to generate these plasmids are listed in the Supplementary Methods. All plasmids were verified by DNA sequencing (Azenta).

### Cell culture

HEK293T, THP-1, and HCC1806 cells were purchased from ATCC. HEK293T and HCC1806 cells were cultured in Dulbecco’s Modified Eagle Medium (DMEM) supplemented with L-glutamine and 10% fetal bovine serum (FBS). THP-1 cells were cultured in RPMI 1640 supplemented with L-glutamine and 10% FBS and differentiated into macrophages with phorbol 12-myristate 13-acetate (PMA, 50 ng/mL) for 48 h. Cells were maintained at 37 °C in a humidified incubator with 5% CO_2_. For hypoxia experiments, cells were maintained at 1% O_2_ and 5% CO_2_ at 37 °C. Cell lines were routinely tested and confirmed to be mycoplasma free.

### Transient transfections

HEK293T or HCC1806 cells were seeded in 96-well culture plates at 50,000 or 25,000 cells per well, respectively, in DMEM supplemented with 10% FBS. The following day, DNA plasmids were diluted in Opti-MEM and transfected using FuGENE HD according to the manufacturer’s instructions.

### Generation of stable cell lines

Lentivirus was generated in HEK293T cells by co-transfection of the indicated expression vector with psPAX2 and pMD2.G using FuGENE HD. For transduction of HEK293T cells, 2 μg expression vector, 2 μg psPAX2, and 1 μg pMD2.G were transfected. Viral supernatants were collected after 48 h, filtered through a 0.45-μm filter, and used to generate cells stably expressing GSDMD, CASP1, and the indicated inducible DIPTAR constructs.

For THP-1 and HCC1806 cells, lentivirus was generated using 10 μg expression vector, 10 μg psPAX2, and 5 μg pMD2.G. Viral supernatants were collected after 48 h, filtered, and concentrated using a PEG virus precipitation kit (Abcam). THP-1 or HCC1806 cells were transduced and after 48 h, cells were selected with hygromycin (100 μg/mL), blasticidin (10 μg/mL), or puromycin (1 μg/mL), as appropriate.

HCC1806 cells stably expressing Cas9-P2A-mCherry were generated using the PiggyBac system. Cells were seeded in 6-well plates at 0.5 x 10^6^ cells per well and co-transfected with Super PiggyBac Transposase (1 μg) and PiggyBac Cas9-P2A-mCherry (0.5 μg). A control lacking transposase was included. After 48 h, cells were selected with blasticidin (10 μg/mL) until all control cells died, followed by fluorescence-activated cell sorting to isolate the top 5% of mCherry-expressing cells.

### siRNA transfections

HCC1806 cells were reverse-transfected with siRNAs (IDT) at a final concentration of 20 μM using Lipofectamine RNAiMAX (Thermo Fisher Scientific) according to the manufacturer’s instructions. Cells were seeded in 96-well plates at 40,000 cells per well. After approximately 16 h, the medium was replaced to minimize transfection-associated cytotoxicity before subsequent experiments. siRNA sequences and target genes are provided in the Supplementary Methods.

### mRNA synthesis and transfection

Codon-optimized ZTA and eGFP sequences were cloned into plasmids containing optimized 5′ and 3′ untranslated regions (UTRs) and a 101-nucleotide poly(A) tail. Plasmids were linearized by restriction digestion and used as templates for in vitro transcription (IVT) with the MEGAscript T7 Transcription Kit (Thermo Fisher Scientific). mRNAs were synthesized with N1-methylpseudouridine-modified nucleosides and co-transcriptionally capped using CleanCap technology (TriLink BioTechnologies). Following IVT, mRNAs were purified by cellulose chromatography to remove double-stranded RNA contaminants, ethanol precipitated and resuspended in nuclease-free water. mRNA integrity was assessed by gel electrophoresis, and innate immune stimulatory activity was evaluated by IFN-α secretion from human monocyte-derived dendritic cells. Purified mRNAs were stored at -80 °C until use.

HCC1806 cells were seeded in 96-well plates at 25,000 cells per well in DMEM containing 10% FBS and transfected with 1 μg of the indicated mRNA using jetMESSENGER mRNA Transfection Reagent (Sartorius, 101000005) according to the manufacturer’s instructions. Cells were incubated for 6 h at 37 °C and 5% CO_2_ before initiating Incucyte measurements.

### Incucyte SYTOX Green and propidium iodide uptake assays

Cell-death kinetics were measured using an Incucyte S3 Live-Cell Analysis System (Sartorius). HEK293T cells were seeded in black, clear-bottom 96-well plates at 50,000 cells per well in DMEM or Opti-MEM containing SYTOX Green (0.2 μM). THP-1 cells were seeded at 80,000 cells per well in RPMI containing PMA (50 ng/mL) and differentiated for 48 h, after which the medium was replaced with Opti-MEM containing SYTOX Green (0.2 μM). HCC1806 cells were seeded at 25,000 cells per well in DMEM, and the medium was replaced the following day with Opti-MEM containing SYTOX Green (0.2 μM) or PI (1 μg/mL). Assays were performed in 100 μL per well. Images were acquired with a 10x objective at 1-h intervals for 6–24 h. SYTOX Green-positive cells were quantified using Incucyte analysis software, and cell death was normalized to cells treated with 0.1% Triton X-100.

### HRE-GFP reporter assays

HCC1806 HRE-GFP reporter cells were transfected with siRNAs as described above. Following the medium change, GFP fluorescence was monitored using an Incucyte S3 Live-Cell Analysis System (Sartorius). GFP-positive objects were quantified and normalized to phase confluence using the Incucyte analysis software. Values are reported as fold change relative to cells transfected with non-targeting control siRNA (siNTC).

### Immunoblotting

Samples were collected as cell lysates, culture supernatants, or combined lysate and supernatant fractions (total). Lysates were collected directly in 2x sample loading buffer or normalized for protein concentration using the DC Protein Assay Kit (Bio-Rad) before addition of sample buffer. Proteins from Opti-MEM supernatants were concentrated by methanol-chloroform precipitation. For total samples, cells and supernatants were collected together, sonicated to lyse cells, and subjected to methanol-chloroform precipitation. Samples were boiled at 95 °C for 10 min before immunoblot analysis.

### FLAG immunoprecipitation

HEK293T cells were seeded in 6-well plates at 0.5 x 10^6^ cells per well and transfected 24 h later with the indicated 2xFLAG-tagged constructs using FuGENE HD. HIF-1α WT, DIPTAR-HIF-1α(FL), DIPTAR-HIF-1α(CT), DIPTAR-HIF-1α(ODD), and DIPTAR-HIF-1α(ODD-P402A/P564A) were transfected at 0.5 μg per well; DIPTAR-HIF-1α(CODD) at 0.2 μg per well; and CARD8 WT, DIPTAR-eGFP, DIPTAR-HIF-1α(CTAD), and GFP at 0.1 μg per well. RFP plasmid was added to a total of 2 μg of DNA per well.

After 48 h, cells were harvested, lysed by sonication, and clarified by centrifugation at 400 x g for 10 min. Protein concentrations were determined using the DC Protein Assay (Bio-Rad). Equal amounts of protein were retained as input or incubated with 100 μL anti-FLAG M2 agarose resin (Sigma-Aldrich) in Pierce Micro-Spin Columns (Thermo Scientific) with end-over-end rotation overnight at 4 °C. The resin was washed three times with 350 μL PBS, and bound proteins were eluted with 100 μL PBS containing 300 ng/μL 3xFLAG peptide (Sigma-Aldrich) for 1 h at room temperature. Input and eluate samples were mixed 1:1 with 2x sample loading buffer, boiled at 95 °C for 10 min, and analyzed by immunoblotting.

### Western blotting

Protein samples were resolved on NuPAGE 4–12% Bis-Tris Midi Protein Gels (Thermo Fisher Scientific, WG1403BOX) at 175 V and transferred to 0.45-μm nitrocellulose membranes (Bio-Rad, 1704271) at 25 V for 7 min using a Trans-Blot Turbo System (Bio-Rad). Membranes were blocked with Intercept Blocking Buffer (LI-COR Biosciences, 927-70010) for 1 h and incubated overnight at 4 °C with primary antibodies diluted 1:1,000 in a 1:1 mixture of blocking buffer and TBS containing 0.1% Tween-20. After three washes, membranes were incubated for 1 h at room temperature with IRDye 800CW- or IRDye 680CW-conjugated donkey anti-mouse, anti-rabbit, or anti-goat secondary antibodies. Membranes were washed three times and imaged using an Odyssey M Imaging System (LI-COR Biosciences). Images were analyzed using Empiria Studio v3.3 (LI-COR Biosciences), and brightness, contrast, and tone were adjusted uniformly across each image.

For densitometric analysis of HIF-1α stability, HIF-1α band intensity was normalized to the corresponding GAPDH loading control and expressed as fold change relative to the non-targeting siRNA control (siNTC).

### Cloning of the Bison library into the DIPTAR CRISPR screening vector

The focused Bison CRISPR library plasmid pool (Addgene) was PCR-amplified using primers containing Gibson assembly overhangs complementary to the MluI and AvrII sites in the DIPTAR-HIF-1α(ODD) CRISPR screening vector. PCR was performed using KAPA HiFi HotStart DNA Polymerase (Roche) in 25-μL reactions containing 10 ng library template and 0.3 μM of each forward and reverse primer. A no-template control was included. Cycling conditions were 95 °C for 3 min; 10 cycles of 98 °C for 20 s, 65 °C for 15 s, and 72 °C for 15 s; and 72 °C for 1 min. PCR products were resolved on a 1.5% agarose gel and purified using the QIAquick Gel Extraction Kit (Qiagen).

The amplified library was assembled into the MluI/AvrII-digested DIPTAR-HIF-1α(ODD) screening vector using NEBuilder HiFi DNA Assembly (NEB) for 1 h at 50 °C. An assembly lacking the library insert was included as a negative control. Assembled DNA was concentrated by isopropanol precipitation and electroporated into Endura electrocompetent cells (Lucigen) at 1,800 V, 10 μF, and 600 Ω. Following recovery, serial dilutions were plated on LB agar containing carbenicillin (100 μg/mL) to determine library representation, and the remaining cells were expanded overnight in 1 L LB containing carbenicillin (100 μg/mL). Library coverage was calculated to be ∼1200x from colony counts and dilution factors. Plasmid DNA was isolated using the QIAGEN Plasmid Maxi Kit according to the manufacturer’s instructions.

### Analysis of DIPTAR based CRISPR knockout screen

Sequencing reads from all seven samples were processed using Cutadapt (version 5.2) to remove the sequence preceding the sgRNA spacer, with GAAACACCG specified as the 5′ adapter. Trimmed reads were quantified using MAGeCK count (version 0.5.9.4) with the BISON library sequence file as the reference to generate an unnormalized sgRNA count matrix. The sgRNA count matrix was analyzed using MAGeCK maximum-likelihood estimation (MLE) with default median normalization to account for differences in library size across samples. A design matrix specifying time point and Dox treatment was used, with the baseline sample as the reference condition. For each gene, the four targeting sgRNAs were jointly modeled to estimate a gene-level β score and corresponding *P* value. Dox-associated β scores and *P* values were used to evaluate gene enrichment at each time point and visualized as β score versus −log10(*P* value). Further information about the quality control and count data can be found as Supplemental Information. The MAGeCK MLE design matrix was as follows:

| Samples | Baseline | 0 h | 6 h | 24 h | Dox_0 h | Dox_6 h | Dox_24 h |
| --- | --- | --- | --- | --- | --- | --- | --- |
| Baseline | 1 | 0 | 0 | 0 | 0 | 0 | 0 |
| 0 h DMSO | 1 | 1 | 0 | 0 | 0 | 0 | 0 |
| 6 h DMSO | 1 | 0 | 1 | 0 | 0 | 0 | 0 |
| 24 h DMSO | 1 | 0 | 0 | 1 | 0 | 0 | 0 |
| 0 h Dox | 1 | 1 | 0 | 0 | 1 | 0 | 0 |
| 6 h Dox | 1 | 0 | 1 | 0 | 0 | 1 | 0 |
| 24 h Dox | 1 | 0 | 0 | 1 | 0 | 0 | 1 |

### Statistical analysis

Statistical analyses were performed using GraphPad Prism 11. Comparisons between two groups were performed using two-tailed Student’s *t*-tests. Comparisons involving multiple groups or experimental factors were performed using one- or two-way ANOVA, as appropriate, followed by Tukey’s multiple-comparisons test. Statistical significance is indicated as *p < 0.05, **p < 0.01, ***p < 0.001, and ****p < 0.0001. The statistical tests, sample sizes, and additional details for individual experiments are provided in the corresponding figure legends.

## Supporting information

Supplemental CRISPR Screen Analysis Info

## Data Availability

All data supporting the findings of this study, including unprocessed gels, are available within the article and Supplementary Information. Sequencing data have been deposited in the NCBI Gene Expression Omnibus (GEO) under accession number GSE345051. Additional data are available from the corresponding author upon reasonable request.

## Author contributions CRediT

Conceptualization: CYT, PME., Methodology: CYT, PME, Investigation: PME, WY, MK, ABM, RCP, CMB, BM, BMD, Visualization: CYT, PME, Funding acquisition: CYT, Project administration: CYT Data curation: CYT, PME, Formal Analysis: CYT, PME, WY, MK, ABM, RCP, CMB, BM, BMD, Resources: CYT Supervision: CYT, Writing – original draft: CYT, PME, Writing – review & editing: CYT, PME, MK, CMB.

## Acknowledgements

We thank all members of the Taabazuing laboratory for helpful scientific discussions. We thank Mark Boyer, Dr. Sunny Shin, and Dr. Frank Lee for access to the hypoxia chamber. We thank Dr. Luca Busino for plasmids and the Bison Library. We thank Dr. Ophir Shalem and Dr. Junwei Shi for discussions regarding CRISPR screens, and Sixiang Yu from the laboratory of Dr. Junwei Shi for reagents and technical assistance related to CRISPR screens. We thank Jacqueline J. Peng for discussions regarding computational analysis of CRISPR screens. We acknowledge Penn Genomics and Sequencing Core: RRID:SCR_024999 for sequencing the CRISPR screening libraries.

This work was supported by an NIGMS Maximizing Investigator’s Research Award (MIRA; Grant No. 1R35GM155239-01) and Burroughs Wellcome Fund Investigators in the Pathogenesis of Infectious Diseases Award (Grant No. 1541395) to CYT, Martin and Pamela Winter Infectious Disease Fellowship to PME, Penn Provost Postdoctoral Fellow and Burroughs Wellcome Fund (Grant No. 1054907) to CMB.

## Declaration of generative AI and AI-assisted technologies

After drafting the manuscript, the authors used ChatGPT 4 to edit the writing. After using this tool, the authors reviewed and revised as needed and take responsibility for the content of the publication.

