## Supplemental Figures, Source Data, and Methods for "DIPTAR: A synthetic biology platform for functional interrogation of protein degradation"

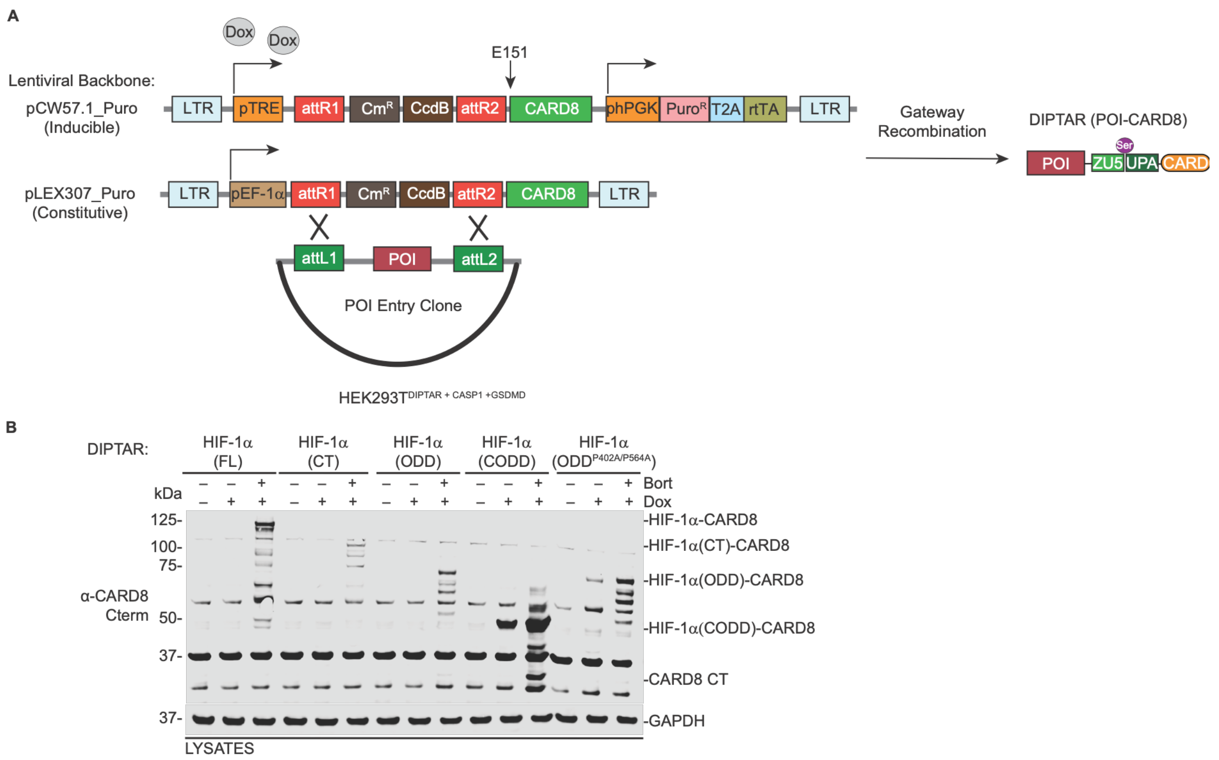


**Fig. S1. Gateway vectors enable modular cloning and expression of POI-CARD8 DIPTARs.** (**A**) Schematic of the DIPTAR Gateway cloning platform with inducible and constitutive vector backbones. (**B**) Immunoblot analysis of HEK293T cells stably expressing the indicated doxycycline-inducible HIF-1α-based DIPTARs. Cells were treated with doxycycline (Dox; 1 μg/mL) and bortezomib (Bort; 1 μM) for 18 h at 21% $O_2$ before lysis.

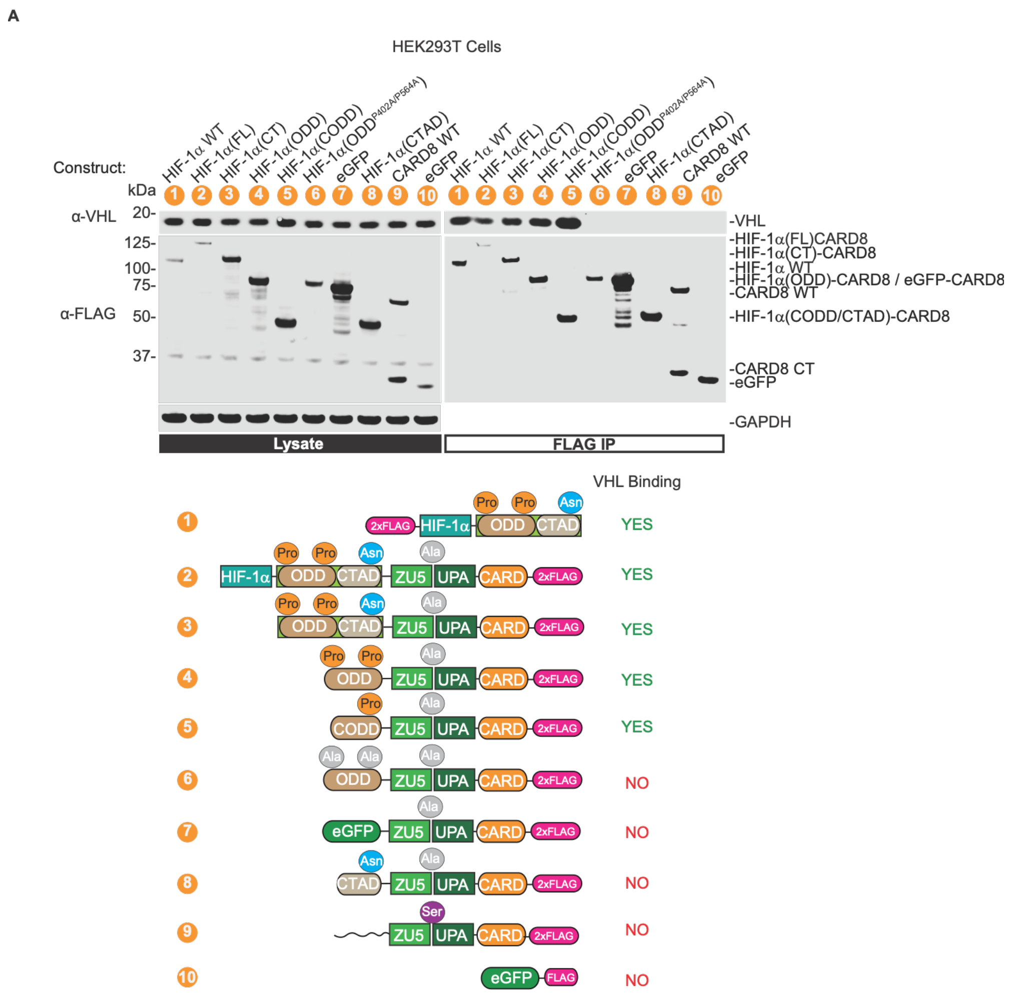


**Fig. S2. DIPTAR-HIF-1α binds VHL in a degron-dependent manner.** (**A**) HEK293T cells were transfected with the indicated FLAG-tagged constructs for 48 h, followed by FLAG immunoprecipitation and immunoblot analysis of endogenous VHL association.

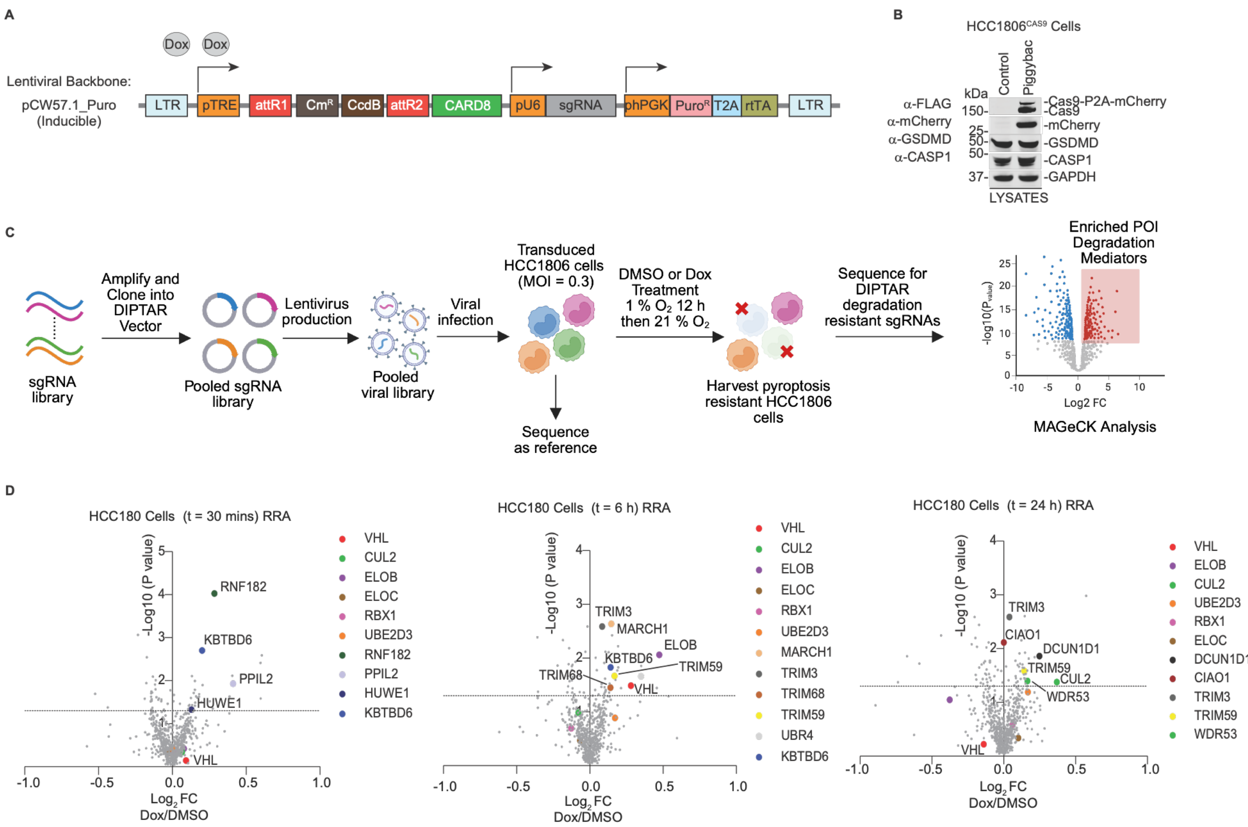


E

| Gene | MLE t=0h Rank | MLE t=6h Rank | MLE t=24h Rank | RRA t=0h Rank | RRA t=6h Rank | RRA t=24h Rank |
|---|---|---|---|---|---|---|
| VHL | - | 6 | - | - | 22 | - |
| CUL2 | - | - | 2 | - | - | 33 |
| ELOB | - | 3 | - | - | 7 | - |
| UBE2D3 | - | - | 5 | - | - | - |
| DCUN1D1 | - | - | 10 | - | - | 9 |
| UBE2M | - | 15 | - | - | - | - |
| HUWE1 | 11 | - | - | 33 | - | - |
| COPS5 | 10 | 2 | - | - | - | - |
| CIAO1 | 8 | 19 | - | - | - | 6 |
| RNF182 | 12 | - | - | 1 | - | - |
| MARCH1 | - | 27 | - | - | 1 | - |
| DDB1 | 6 | - | - | - | - | - |
| DTL | 14 | - | 25 | - | - | - |
| PPIL2 | 4 | 13 | - | 8 | - | - |
| TRIM3 | - | 21 | - | - | 2 | 3 |
| TRIM59 | - | 17 | 17 | - | 16 | 21 |
| TRIM68 | - | 23 | - | - | 25 | - |
| UBR4 | - | 11 | - | - | 17 | - |
| TRIP12 | - | - | 15 | - | - | - |
| KBTBD6 | 18 | - | - | 2 | 13 | - |
| WDR26 | 21 | - | - | - | - | - |
| WDR53 | - | - | 24 | - | - | 31 |

**Fig. S3. DIPTAR-based CRISPR screening identifies regulators of HIF-1α degradation.** (**A**) Schematic of the DIPTAR vector used for pooled CRISPR screening, containing a U6-driven sgRNA expression cassette. (**B**) Immunoblot confirming Cas9 expression in HCC1806 cells. (**C**) Schematic of the DIPTAR-based pooled CRISPR screening workflow. (**D**) Volcano plots showing gene enrichment at the indicated time points by MAGeCK robust rank aggregation (RRA) analysis. (**E**) Comparison of gene rankings and enriched hits across time points identified by MAGeCK maximum likelihood estimation (MLE) and RRA analyses.

### Source Data for Figure 1 and Supplemental Figure 1.

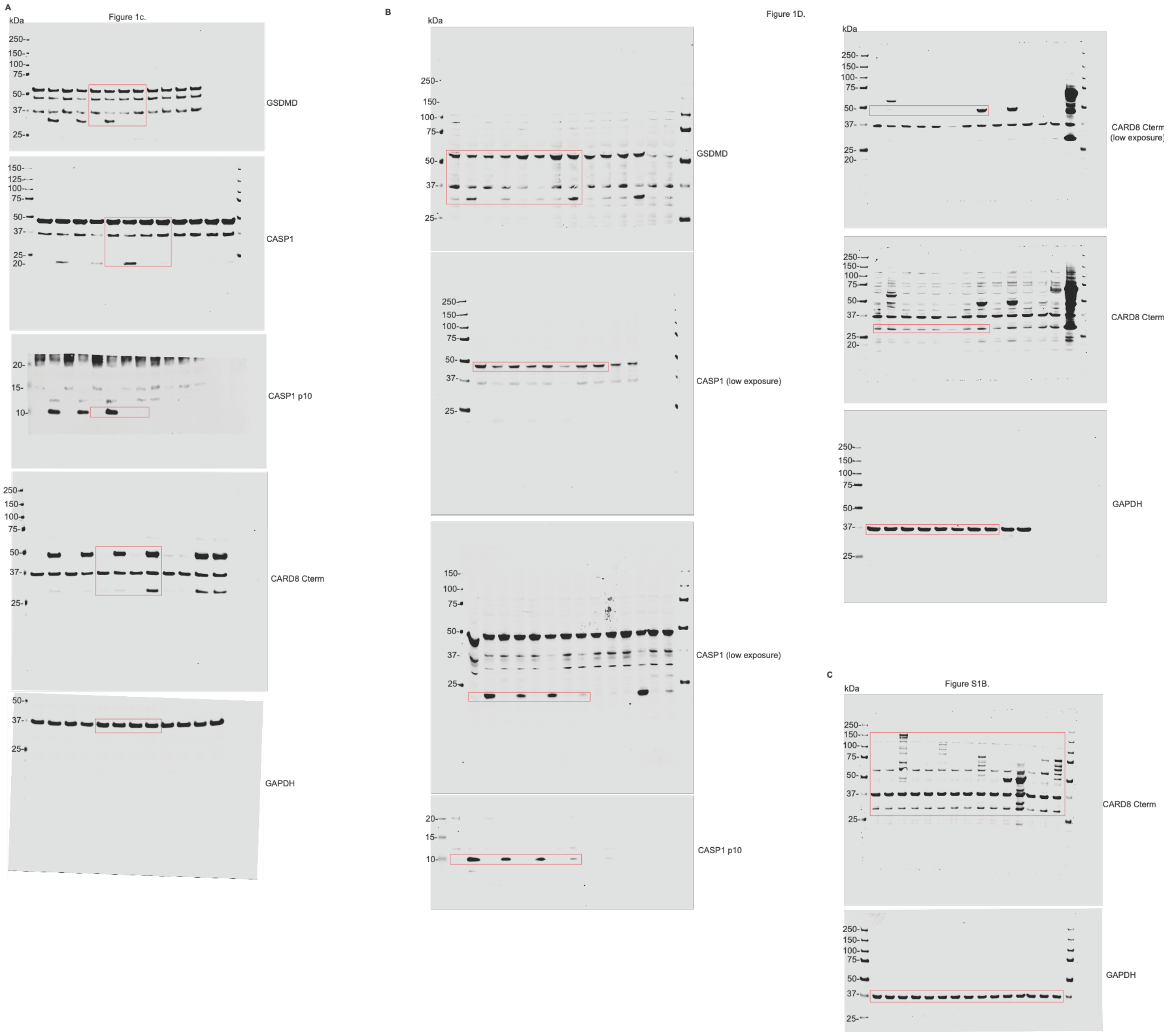

### Source Data for Figure 2 and Supplemental Figure 2.

D

Figure 2a.

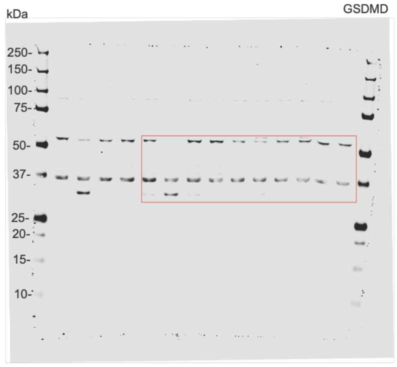


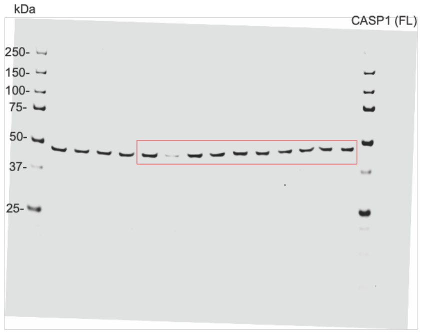


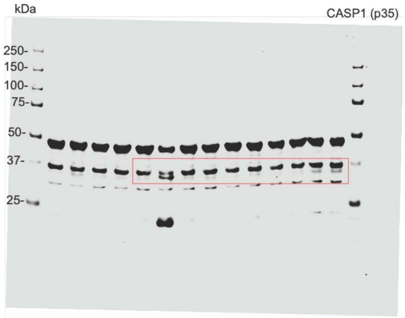


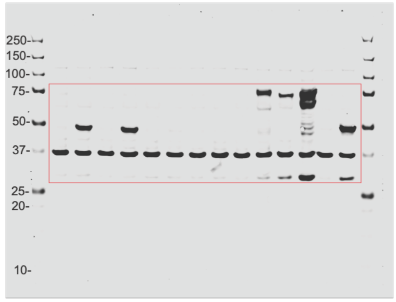


CARD8Cterm

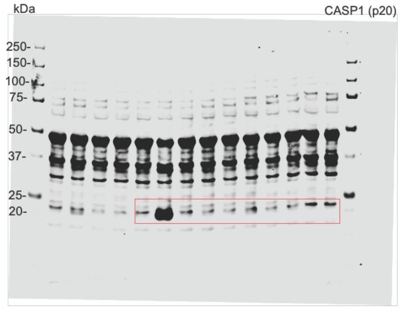


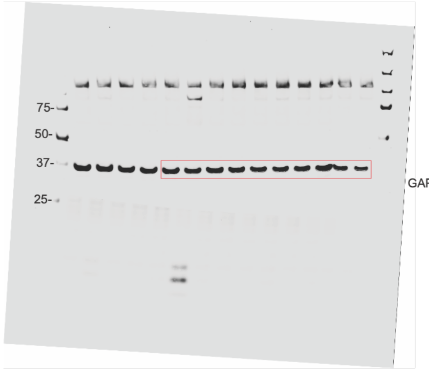


GAPDH

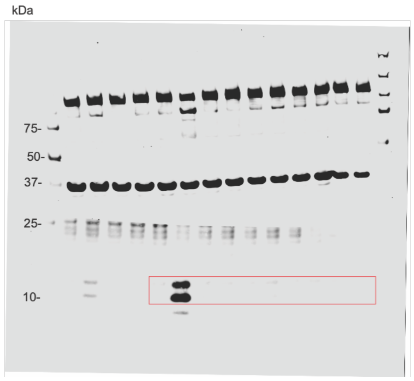


CASP1p10

E

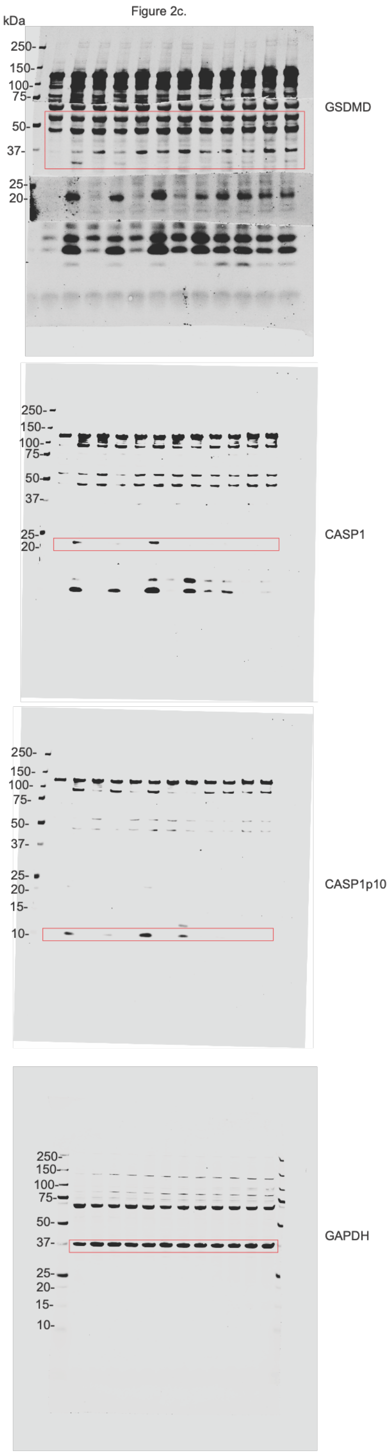
Figure 2c.
kDa
GSDMD
CASP1
CASP1p10
GAPDH

F

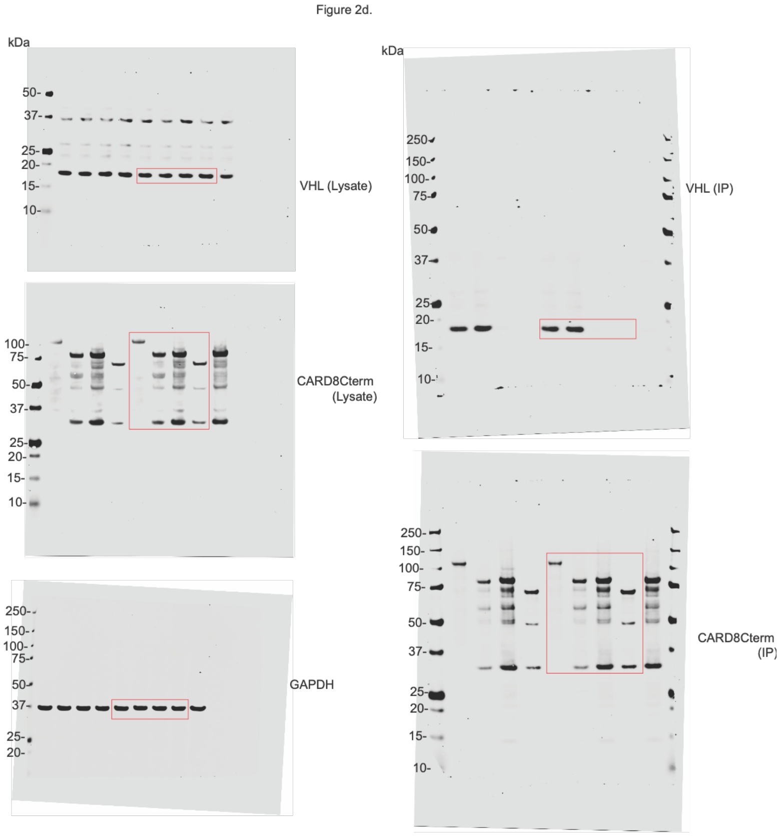
Figure 2d.
kDa
VHL (Lysate)
CARD8Cterm (Lysate)
GAPDH
VHL (IP)
CARD8Cterm (IP)

G

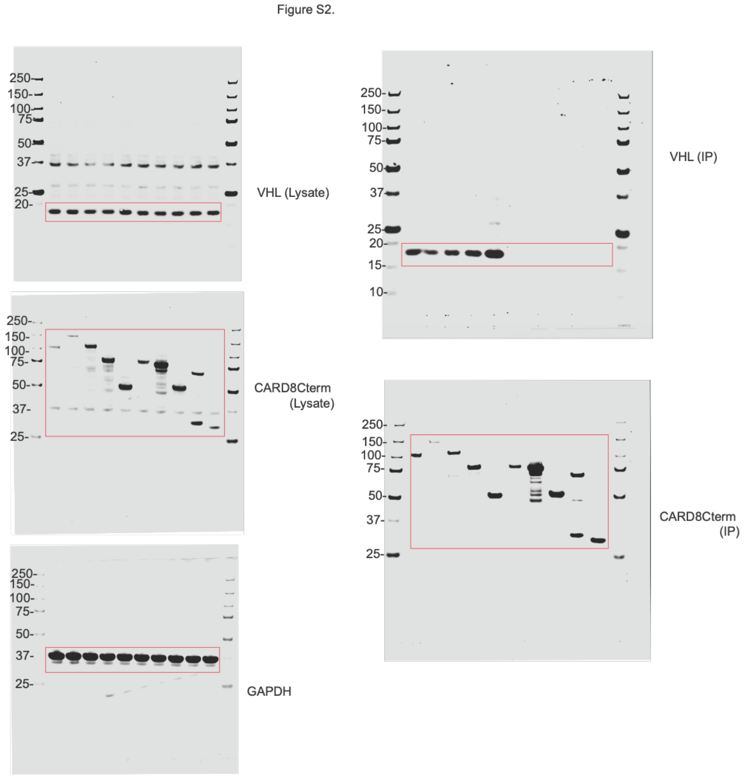
Figure S2.
VHL (Lysate)
CARD8Cterm (Lysate)
GAPDH
VHL (IP)
CARD8Cterm (IP)

### Source Data for Figure 3.

H

Figure 3B.

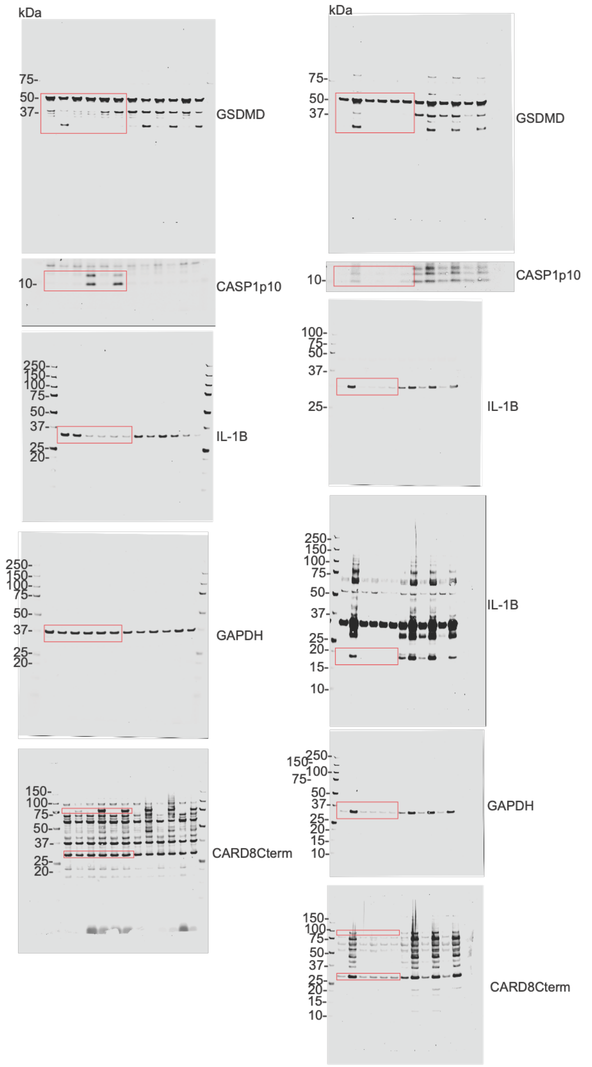


I

Figure 3D.

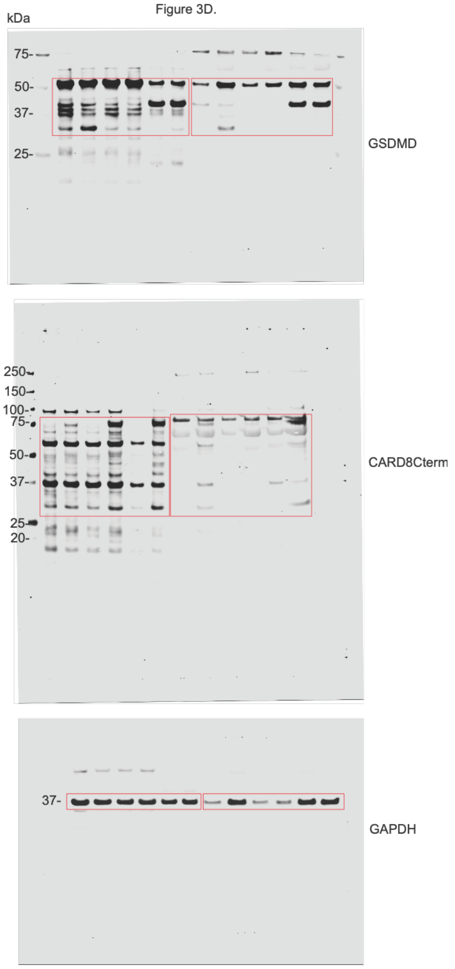


J

Figure 3F.

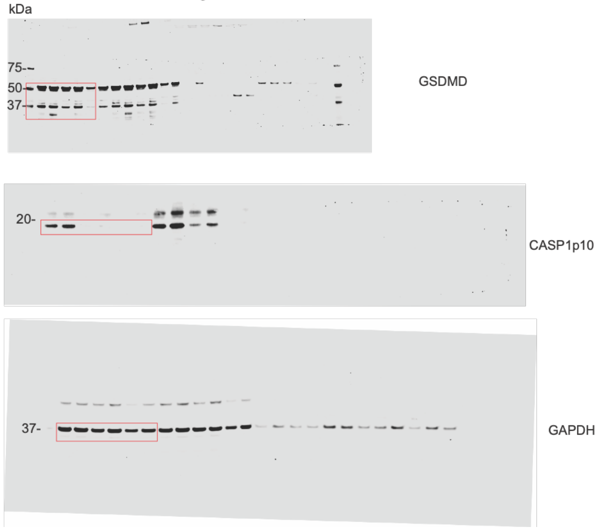

### Source Data for Figure 4 and Supplemental Figure 3.

**K**

Figure S3B.

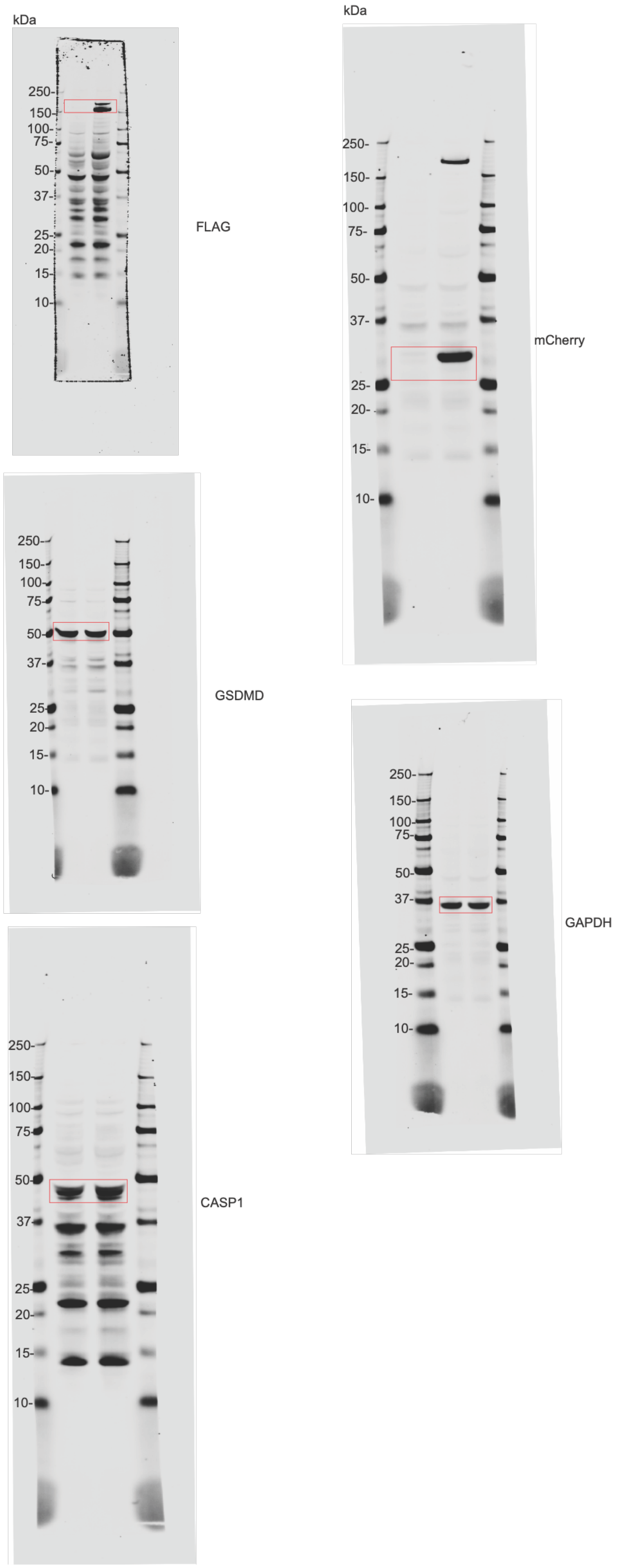


**L**

Figure 4I.

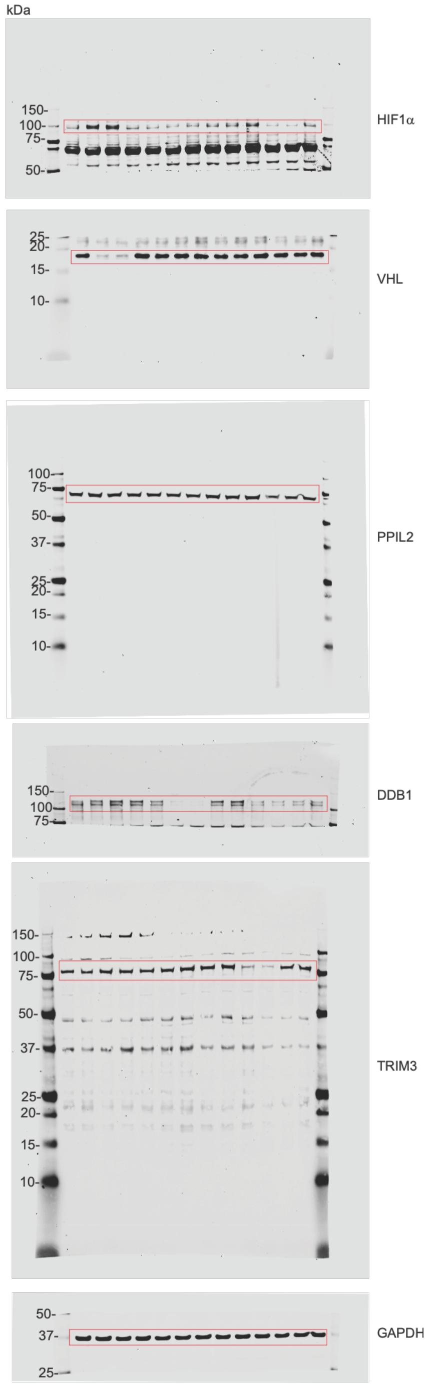

### Source Data for Figure 5.

M

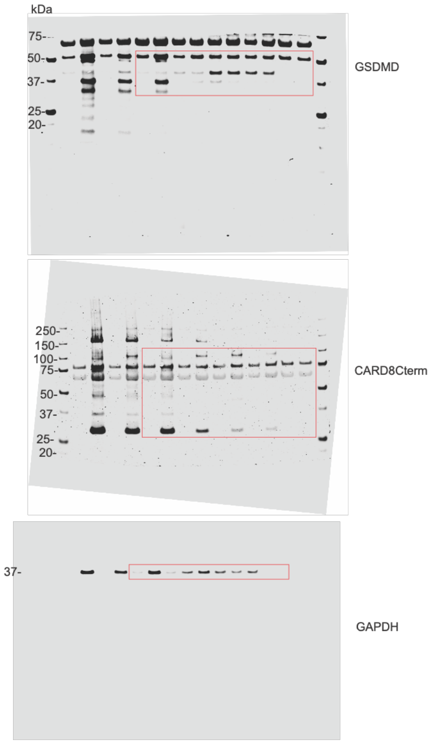


Figure 5C.

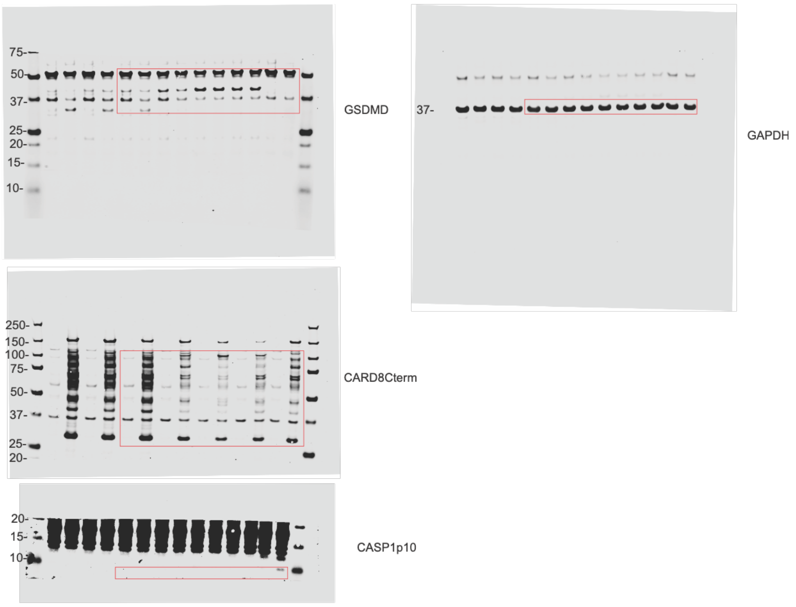


Figure 5E.

N

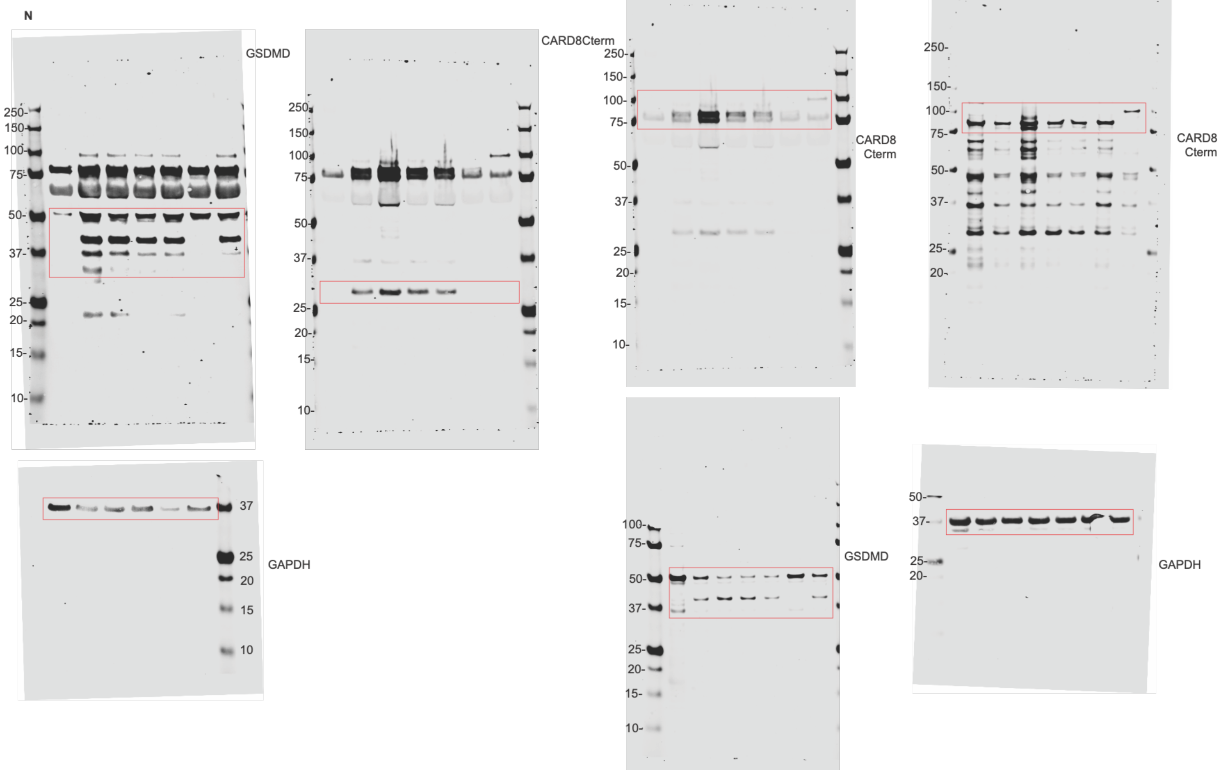

Figure 5G.

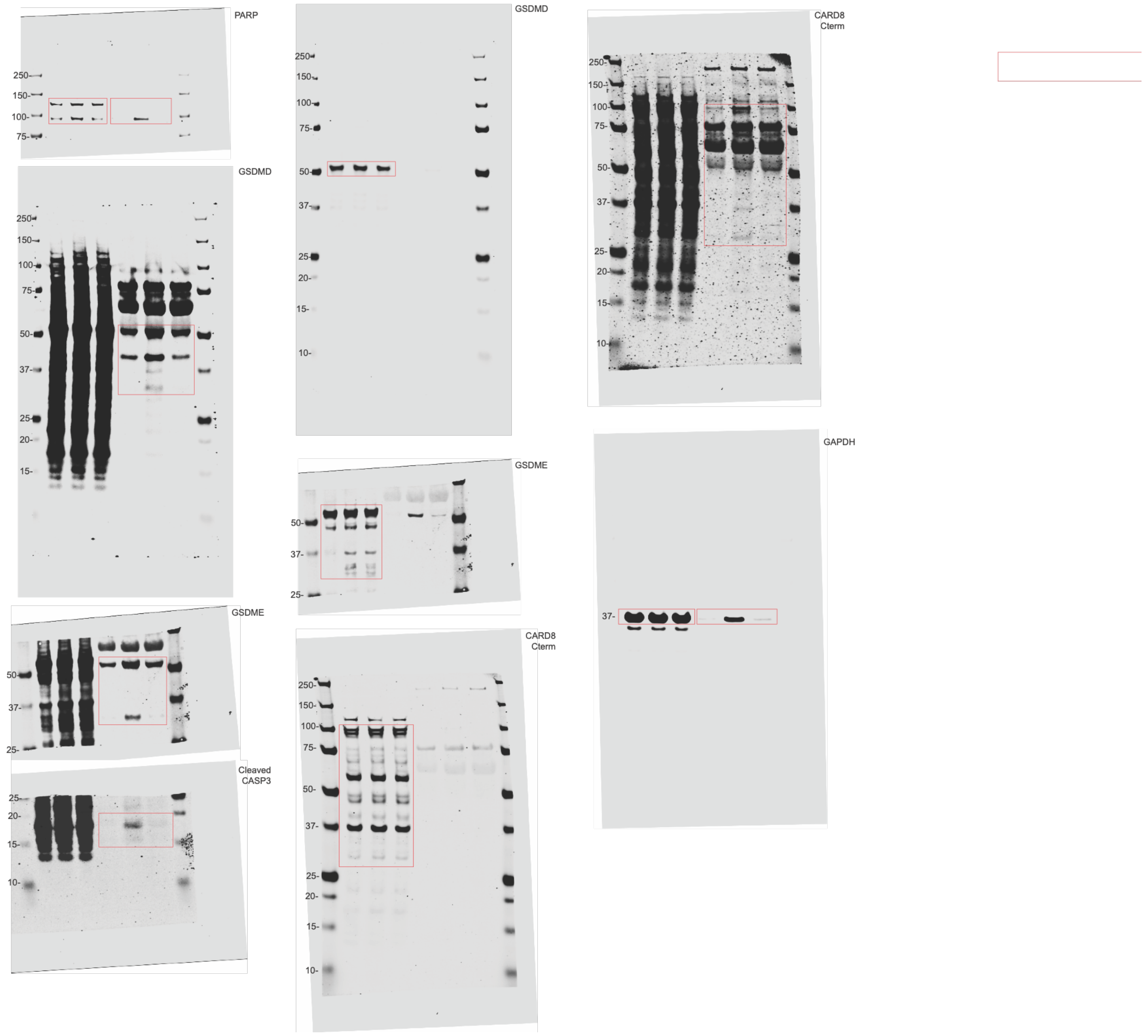

PARP
GSDMD
GSDMD
CARD8 Cterm
GSDME
GAPDH
GSDME
Cleaved CASP3
CARD8 Cterm
250-
150-
100-
75-
50-
37-
25-
20-
15-
10-

### Source Data for Figure 6.

Figure 6B.

P

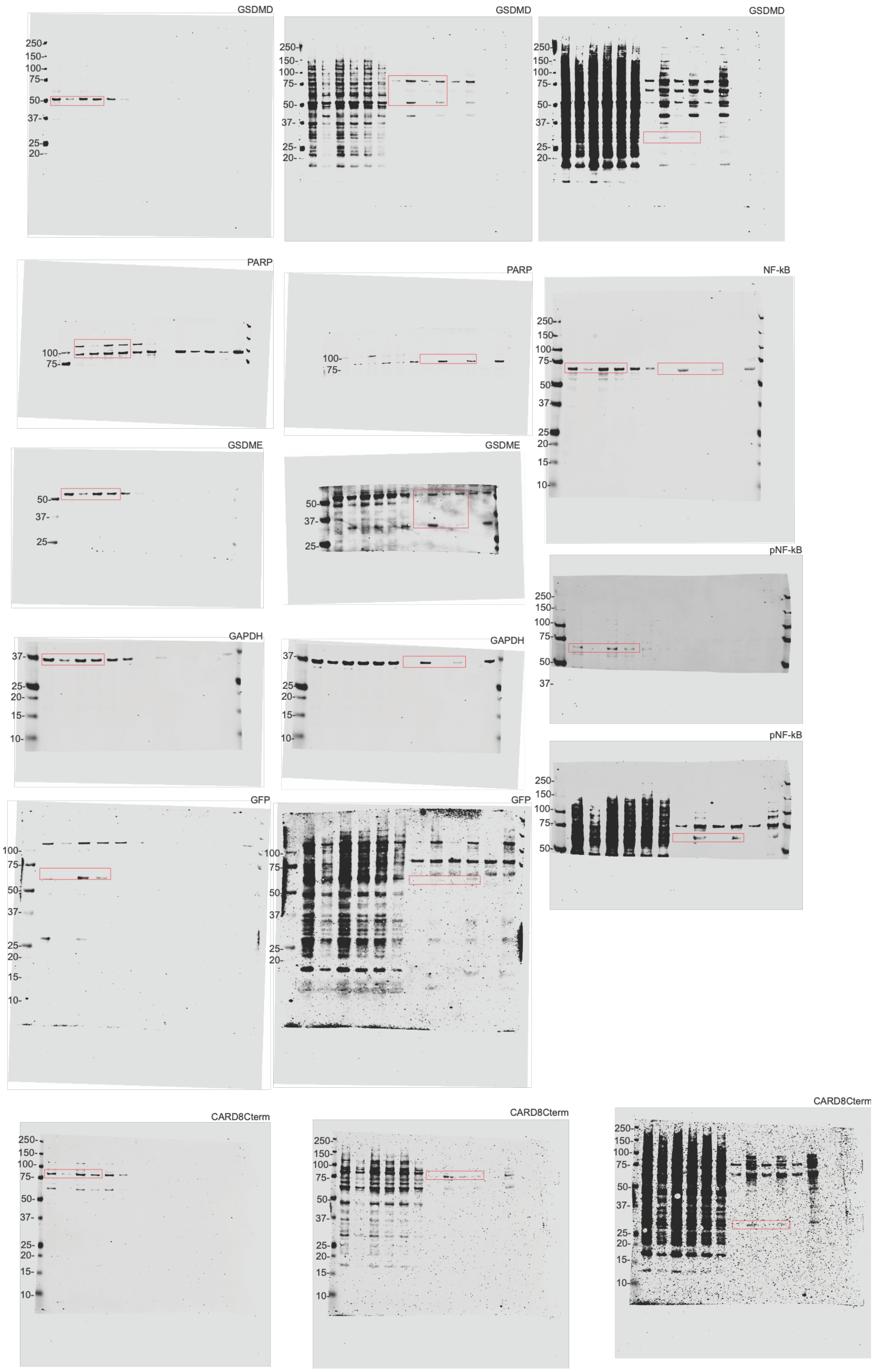

Figure 6D.

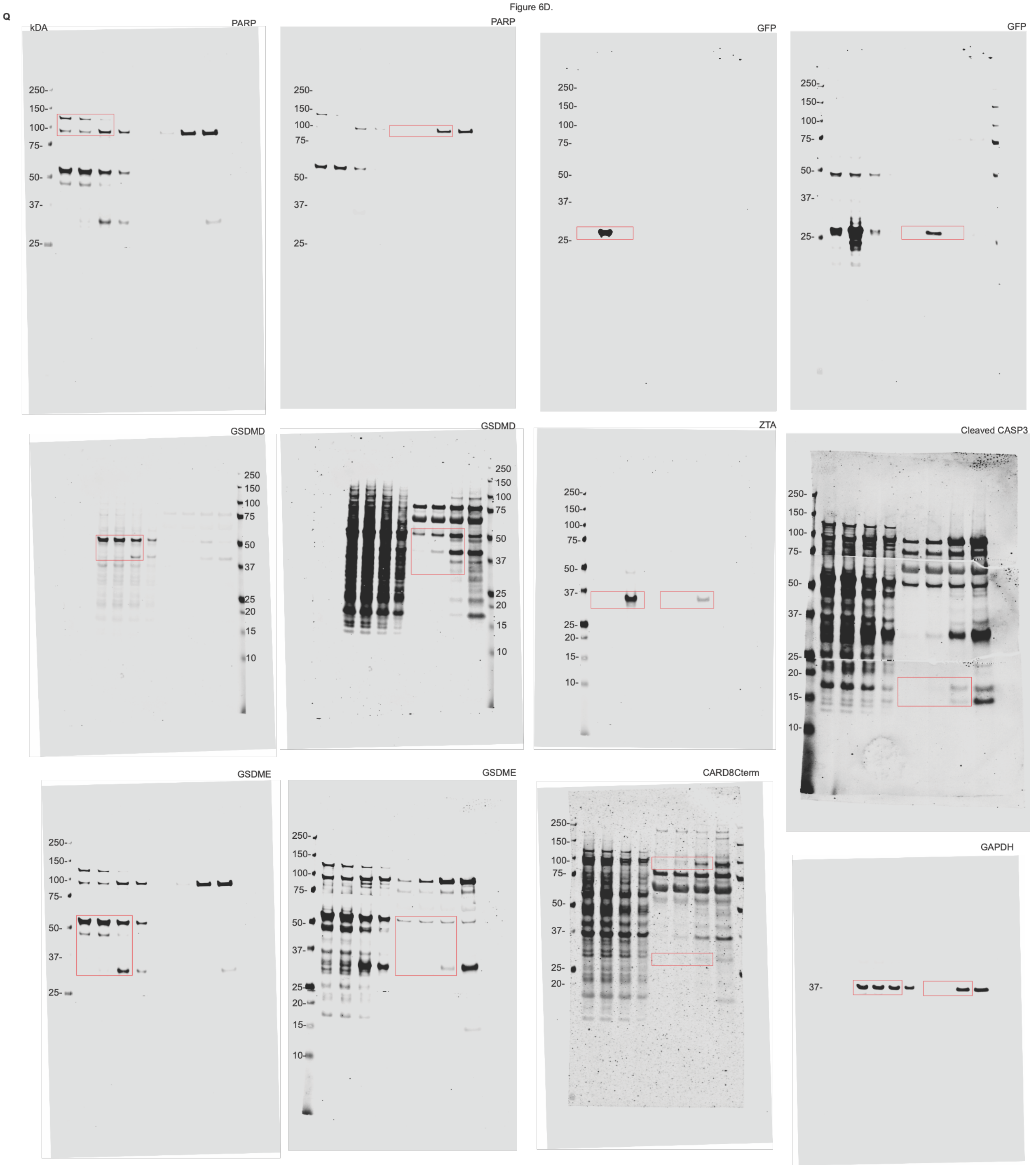
Q
kDA
PARP
PARP
GFP
GFP
GSDMD
GSDMD
ZTA
Cleaved CASP3
GSDME
GSDME
CARD8Cterm
GAPDH

#### Supplementary Methods

##### Supplementary Table 1: Reagents and key resources used in this study

| REAGENT or RESOURCE | SOURCE | IDENTIFIER |
|---|---|---|
| **Antibodies** | | |
| GSDMD Rabbit polyclonal Ab | Novus Biologicals | Cat# NBP2-33422; RRID: AB_2687913 |
| FLAG® M2 Mouse monoclonal Ab | Sigma | Cat# F3165; RRID: AB_259529 |
| GAPDH Rabbit monoclonal Ab | Cell Signaling Tech | Cat# 2118S; RRID: AB_561053 |
| CASP1 p20 Rabbit polyclonal Ab | Cell Signaling Tech | Cat# 2225s; RRID: AB_2243894 |
| CASP1 p12/10 Rabbit monoclonal Ab | Abcam | Cat# ab179515; RRID: AB_2884954 |
| CARD8 Rabbit polyclonal Ab | Abcam | Cat# ab24186 RRID: AB_2275096 |
| hIL-1β Goat Polyclonal Ab | R&D systems | Cat# AF-201-NA; RRID: AB_354387 |
| VHL Rabbit polyclonal Ab | Cell Signaling Tech | Cat# 68547S; RRID: AB_2716279 |
| HIF-1 alpha polyclonal Ab | Proteintech | Cat# 20960-1-AP; RRID: AB_10732601 |
| PARP Rabbit polyclonal Ab | Cell Signaling Tech | Cat# 9542S; RRID: AB_2160739 |
| GSDME Rabbit monoclonal Ab | abcam | Cat# ab215191; RRID: AB_2737000 |
| Cleaved Caspase-3 (Asp175) (5A1E) Rabbit monoclonal Ab | Cell Signaling Tech | Cat# 9664S; RRID: AB_2737000 |
| eGFP Mouse monoclonal Ab | ThermoFisher | Cat# MA1-952; RRID: AB_889471 |
| EBV ZEBRA (BZ1) | Santa Cruz Biotechnology | Cat# sc-53904; RRID: AB_783257 |
| Phospho-NF-κB p65 (Ser536) (93H1) Rabbit monoclonal Ab | Cell Signaling Tech | Cat# 3033S; RRID: AB_331284 |
| NF-kappaB p65 (D14E12) Rabbit monoclonal Ab | Cell Signaling Tech | Cat# 8242S; RRID: AB_10859369 |
| PPIL2 Recombinant Monoclonal Ab | Proteintech | Cat# 87496-1-RR; RRID: AB_3745581 |
| DDB1 Polyclonal Ab | Proteintech | Cat# 11380-1-AP; RRID: AB_2088808 |
| TRIM3 Polyclonal Ab | Proteintech | Cat# 28392-1-AP; RRID: AB_2881131 |
| IRDye 800CW anti-rabbit | LI-COR | Cat# 926-32213; RRID: AB_621848 |

| | | |
|---|---|---|
| IRDye 800CW anti-mouse | LI-COR | Cat# 926-32212;<br>RRID: AB_621847 |
| IRDye 800CW anti-goat | LI-COR | Cat# 926-32214;<br>RRID: AB_621846 |
| IRDye 680CW anti-rabbit | LI-COR | Cat# 926-68073;<br>RRID: AB_10954442 |
| IRDye 680CW anti-mouse | LI-COR | Cat# 926-68072;<br>RRID: AB_10953628 |
| IRDye 680CW anti-goat | LI-COR | Cat# 926-68074;<br>RRID: AB_10956736 |
| **Chemicals, peptides, and recombinant proteins** | | |
| Doxycycline hydrochloride | Fisher Scientific | Cat#ICN19504401 |
| VX-765 | Apexbio Technology LLC | Cat#50-101-3604 |
| Z-VAD-FMK | Enzo Life Sciences | Cat#NC9471015 |
| MLN4924 | Fisher Scientific | Cat#50-187-3933 |
| Bortezomib | LC Laboratories | Cat#B-1408 |
| dBET1 | MedChemExpress | Cat#HY-101838 |
| EM12-SO2F | MedChemExpress | Cat#HY-W760183 |
| GS143 | MedChemExpress | Cat#HY-110261 |
| Recombinant human TNF-α | Invivogen | Cat#rcyc-htnfa |
| FuGENE HD | Promega | Cat#2311 |
| jetMESSENGER® mRNA Transfection | Sartorius | Cat#101000056 |
| ANTI-FLAG® M2 Affinity Gel | Neta Scientific | Cat#A2220 |
| 3xFLAG™ Peptide | Sigma-Aldrich | Cat#F4799 |
| phorbol 12-myristate 12-acetate | Promega | Cat#V1171 |
| **Critical commercial assays** | | |
| DC Protein Assay kit | Bio-Rad | Cat#5000111 |
| SYTOX™ Green Nucleic Acid Stain | Fisher Scientific | Cat#S7020 |
| Propidium Iodide | Cayman Chemical Company | Cat#10008351 |
| Qiagen QIAprep Spin Miniprep Kit | QIAGEN | Cat#27106 |
| QIAGEN Plasmid Plus Midi Kit | QIAGEN | Cat#12943 |
| QIAGEN Plasmid Maxi Kit | QIAGEN | Cat#12162 |
| QIAamp DNA Blood Midi Kit | QIAGEN | Cat#51183 |
| Endura™ Electrocompetent Cells | Lucigen | Cat#60242-1 |
| **Experimental models: Cell lines** | | |
| HEK293T | ATCC | N/A |
| HEK293T CASP1/GSDMD | This Study | N/A |
| HEK293T CASP1/GSDMD/pCW57.1_HIF1α(FL)-CARD8 | This Study | N/A |
| HEK293T CASP1/GSDMD/pCW57.1_HIF1α(CT)-CARD8 | This Study | N/A |
| HEK293T CASP1/GSDMD/pCW57.1_HIF1α(ODD)-CARD8 | This Study | N/A |
| HEK293T CASP1/GSDMD/pCW57.1_HIF1α(ODD)-CARD8[S297A] | This Study | N/A |

| | | |
|---|---|---|
| HEK293T CASP1/GSDMD/pCW57.1_HIF1α(ODD[P402A/P564A])-CARD8 | This Study | N/A |
| HEK293T CASP1/GSDMD/pCW57.1_HIF1α(CODD)-CARD8 | This Study | N/A |
| HEK293T CASP1/GSDMD/pCW57.1_eGFP-CARD8 | This Study | N/A |
| HEK293T CASP1/GSDMD/pCW57.1_ HIF1α(CTAD)-CARD8 | This Study | N/A |
| THP-1 pCW57.1_HIF1α(ODD)-CARD8 | This Study | N/A |
| HCC1806 pCW57.1_HIF1α(ODD)-CARD8 | This Study | N/A |
| HCC1806 CAS9-NLS-FLAG-P2A-mcherry | This Study | N/A |
| HCC1806 HRE-GFP | This Study | N/A |
| HCC1806 pCW57.1_BRD4-CARD8 | This Study | N/A |
| HCC1806 pLEX307_Iκβα-CARD8 | This Study | N/A |
| HCC1806 pLEX307_P53-CARD8 | This Study | N/A |
| **Oligonucleotides** | | |
| pCW57.1_DIPTAR F:<br>5'- ttcagctttcttgtacaaagtggttaccggt gagatcgaagaagattataaaaatcgtca-3' | CARD8/pCW57.1 overlap | N/A |
| pCW57.1_DIPTAR R:<br>5'- tccttagtggtggtggtggtggtggaccggttcattacta caaattctgctgtctaagataggacacg-3' | CARD8/pCW57.1 overlap | N/A |
| pLEX_307_DIPTAR F:<br>ttcagctttcttgtacaaagtggtT GATATC gagatcgaagaagattataaaaatcgtca | CARD8/pLEX307 overlap | N/A |
| pLEX_307_DIPTAR R:<br>cgcgttgattgtcgacttaaccggtactagttcattacta caaattctgctgtctaagataggacacg | CARD8/pLEX307 overlap | N/A |
| pLEX_307_DIPTAR_2XFLAG R:<br>cgcgttgattgtcgacttaaccggtactagttcattacta cttgtcgtcatcgtccttgtagtccttgtcgtcatcgtccttgtagtcgccgcc caaattctgctgtctaagataggacacg | CARD8/pLEX307 overlap (2XFLAG) | N/A |
| pCW57.1_DIPTAR_U6_sgRNA piece 1 R:<br>tctgctgtccctgtaataaacccgaaccggTtcattactacaaattctgctgtc | CARD8/pCW57.1 overlap/U6-sgRNA cassette | N/A |
| pCW57.1_DIPTAR_U6_sgRNA piece 2 F:<br>tcgggtttattacagggacagcagagatccagtttggttaattaaggtaccgagg g | CARD8/pCW57.1 overlap/U6-sgRNA cassette | N/A |
| pCW57.1_DIPTAR_U6_sgRNA piece 2 R:<br>tgtattttgtgtgctcatttacaacACGCGTttcgtcctttccacaagatatata aagccaagaaatcg | CARD8/pCW57.1 overlap/U6-sgRNA cassette | N/A |
| pCW57.1_DIPTAR_U6_sgRNA piece 3 F:<br>tatatatcttgtggaaaggacgaaACGCGTgttgtaaatgagcacacaaa atacacatgc | CARD8/pCW57.1 overlap/U6-sgRNA cassette | N/A |
| pCW57.1_DIPTAR_U6_sgRNA piece 3 R:<br>taacttgctatttctagctctaaaaCCTAGGtacaaaaaagagcaagaag c | CARD8/pCW57.1 overlap/U6-sgRNA cassette | N/A |

| | | |
|---|---|---|
| pCW57.1_DIPTAR_U6_sgRNA piece 4 F: ttttagcttcttgctcttttttgtaCCTAGGttttagagctagaaatagcaagttaaaataaggctagtcc | CARD8/pCW57.1 overlap/U6-sgRNA cassette | N/A |
| pCW57.1_DIPTAR_U6_sgRNA piece 4 R: cttagtggtggtggtggtggaccggactgacgggcaccggagc | CARD8/pCW57.1 overlap/U6-sgRNA cassette | N/A |
| hCARD8 S297A Quick Change F: ctggaaagccccagcttcgccctgatgggcatcctgctgc | | N/A |
| hCARD8 S297A Quick Change R: gcagcaggatgcccatcagggcgaagctggggctttccag | | N/A |
| hCARD8 S297A Q5 SDM F: cttcgccctgatgggcatcctgc | | N/A |
| hCARD8 S297A Q5 SDM R: ctggggctttccaggacagc | | N/A |
| HIF1α(FL) attb F: GGGGACAAGTTTGTACAAAAAAGCAGGCTTCGCC atg gagggcgccggcggcgcg | For pDONR221 | N/A |
| HIF1α(CT) / HIF1α(ODD) attb F: GGGGACAAGTTTGTACAAAAAAGCAGGCTTCGCC ATGgccccagccgctggagacaca | For pDONR221 | N/A |
| HIF1α(FL) / HIF1α(CT) attb R: GGGGACCACTTTGTACAAGAAAGCTGGGTC gttaacttgatccaaagctctgagtaattc | For pDONR221 | |
| HIF1α(ODD) attb R: GGGGACCACTTTGTACAAGAAAGCTGGGTC ctggaatactgtaactgtgctttgaggact | For pDONR221 | N/A |
| HIF1α(CODD) attb F: GTACAAAAAAGCAGGCTTCGCCACC ATGgatttagacttggagatgttagctccctatatcccaatggatgatgacttccagtta | For pDONR221 | N/A |
| HIF1α(CODD) attb R: GGGGACCACTTTGTACAAGAAAGCTGGGTC taactggaagtcatcatccattgggatatagggagctaacatctccaagtctaaatc | For pDONR221 | N/A |
| HIF1α(CTAD) attb F: GTACAAAAAAGCAGGCTTCGCCACCATG gatgaaagtggattaccacagctgacc | For pDONR221 | N/A |
| HIF1α(CTAD) attb R: GGGGACCACTTTGTACAAGAAAGCTGGGTCtataggagcattaacttcacaatcataact | For pDONR221 | N/A |
| eGFP attb F: GGGGACAAGTTTGTACAAAAAAGCAGGCTTCGCC atggtgagcaagggcgaggagctg | For pDONR221 | N/A |
| eGFP attb R: GGGGACCACTTTGTACAAGAAAGCTGGGTC CTTGTACAGCTCGTCCAT | For pDONR221 | N/A |
| HIF1α(ODD) P402A Q5 SDM F: CGCCATGGCCgcgGCCGCTGGAG | | N/A |

| | | |
|---|---|---|
| HIF1α(ODD) P402A Q5 SDM R: AAGCCTGCTTTTTTGTACAAACTTGTTGATGCTAGCC | | N/A |
| HIF1α(ODD) P564A Q5 SDM F GATGTTAGCTgcgTATATCCCAATG | | N/A |
| HIF1α(ODD) P564A Q5SDM R: TCCAAGTCTAAATCTGTG | | N/A |
| DN-CUL2 attB F: GTACAAAAAAGCAGGCTTCGCCACC atgtctttgaaaccaagagtagtagattttgatgaaacatgg | For pDONR221 | N/A |
| DN-CUL2 attB R: GGGGACCACTTTGTACAAGAAAGCTGGGTC gaacactgtgatgaagctcgtgagc | For pDONR221 | N/A |
| Iκβα attb F: GGGGACAAGTTTGTACAAAAAAGCAGGCTTCGCC atgttccaggcggccgagcgcccc | For pDONR221 | N/A |
| Iκβα attb R: GGGGACCACTTTGTACAAGAAAGCTGGGTC taacgtcagacgctggcctcc | For pDONR221 | N/A |
| P53 attb F: GTACAAAAAAGCAGGCTTCGCCACC atggaggagccgcagtcag | For pDONR221 | N/A |
| P53 attb R: GGGGACCACTTTGTACAAGAAAGCTGGGTC gtctgagtcaggcccttctgtc | For pDONR221 | N/A |
| CRISPR Library Amplify F: ttaaaattatgttttaaaatggactatcatatgcttaccgtaacttgaaagtatttc gatttcttGGCTTTATATATCTTGTGGAAAGGACGAAACA CCG | Bison Library/ pCW57.1_DIPTAR _U6-sgRNA Overlap | N/A |
| CRISPR Library Amplify R: CTAGCCTTATTTTAACTTGCTATTTCTAGCTCTAAAAC | Bison Library/ pCW57.1_DIPTAR _U6-sgRNA Overlap | N/A |
| PCR1_F1: TCGTCGGCAGCGTCAGATGTGTATAAGAGACAG T TTGTGGAAAGGACGAAACACCG | NGS PCR1 Stagger 1 | N/A |
| PCR1_F2: TCGTCGGCAGCGTCAGATGTGTATAAGAGACAG AT TTGTGGAAAGGACGAAACACCG | NGS PCR1 Stagger 2 | N/A |
| PCR1_F3: TCGTCGGCAGCGTCAGATGTGTATAAGAGACAG GAT TTGTGGAAAGGACGAAACACCG | NGS PCR1 Stagger 3 | N/A |
| PCR1_F4: TCGTCGGCAGCGTCAGATGTGTATAAGAGACAG CGAT TTGTGGAAAGGACGAAACACCG | NGS PCR1 Stagger 4 | N/A |
| PCR1_F5: TCGTCGGCAGCGTCAGATGTGTATAAGAGACAG TCGAT TTGTGGAAAGGACGAAACACCG | NGS PCR1 Stagger 5 | N/A |

| | | |
|---|---|---|
| PCR1_F6:<br>TCGTCGGCAGCGTCAGATGTGTATAAGAGACAG<br>ATCGAT TTGTGGAAAGGACGAAACACCG | NGS PCR1<br>Stagger 6 | N/A |
| PCR1_F7:<br>TCGTCGGCAGCGTCAGATGTGTATAAGAGACAG<br>GATCGAT TTGTGGAAAGGACGAAACACCG | NGS PCR1<br>Stagger 7 | N/A |
| PCR1_F8:<br>TCGTCGGCAGCGTCAGATGTGTATAAGAGACAG<br>CGATCGAT TTGTGGAAAGGACGAAACACCG | NGS PCR1<br>Stagger 8 | N/A |
| PCR1_R1:<br>GTCTCGTGGGCTCGGAGATGTGTATAAGAGACAG<br>CCGACTCGGTGCCACTTTTTCAA | NGS PCR1 | N/A |
| PCR2_F1:<br>AATGATACGGCGACCACCGAGATCTACAC<br>TCGTCGGCAGCGTC | NGS PCR2 | N/A |
| PCR2_R8:<br>CAAGCAGAAGACGGCATACGAGAT TACGATCG<br>GTCTCGTGGGCTCGG | NGS PCR2<br>i7: CGATCGTA | N/A |
| PCR2_R9:<br>CAAGCAGAAGACGGCATACGAGAT CGTGTAGC<br>GTCTCGTGGGCTCGG | NGS PCR2<br>i7: GCTACACG | N/A |
| PCR2_R10:<br>CAAGCAGAAGACGGCATACGAGAT ATCGTGAC<br>GTCTCGTGGGCTCGG | NGS PCR2<br>i7: GTCACGAT | N/A |
| PCR2_R11:<br>CAAGCAGAAGACGGCATACGAGAT AGGCTATA<br>GTCTCGTGGGCTCGG | NGS PCR2<br>i7: TATAGCCT | N/A |
| PCR2_R12:<br>CAAGCAGAAGACGGCATACGAGAT TCTGAGTC<br>GTCTCGTGGGCTCGG | NGS PCR2<br>i7: GACTCAGA | N/A |
| PCR2_R13:<br>CAAGCAGAAGACGGCATACGAGAT CAGGACGT<br>GTCTCGTGGGCTCGG | NGS PCR2<br>i7: ACGTCCTG | N/A |
| PCR2_R14:<br>CAAGCAGAAGACGGCATACGAGAT GTACTGAC<br>GTCTCGTGGGCTCGG | NGS PCR2<br>i7: GTCAGTAC | N/A |
| **siRNA** | | N/A |
| NC-1 siRNA Negative Control:<br>(Proprietary) | IDT | N/A |
| hs.Ri.VHL.13.1<br>Top strand (5′→3′):<br>GAUUUCUGUUGAAACUUACACUGTT<br>Bottom strand (5′→3′):<br>AACAGUGUAAGUUUCAACAGAAAUCUU | IDT | N/A |

| | | |
|---|---|---|
| hs.Ri.VHL.13.2<br>Top strand (5′→3′):<br>GUCAUCUUCUGCAAUCGCAGUCCGC<br>Bottom strand (5′→3′):<br>GCGGACUGCGAUUGCAGAAGAUGACCU | IDT | N/A |
| hs.Ri.PPIL2.13.1<br>Top strand (5′→3′):<br>GUGGAACAGCUAAAUAUCAAGGCCA<br>Bottom strand (5′→3′):<br>UGGCCUUGAUAUUUAGCUGUUCCACUG | IDT | N/A |
| hs.Ri.PPIL2.13.2<br>Top strand (5′→3′):<br>CGUCUUUGACUUACUGAACAUUGTT<br>Bottom strand (5′→3′):<br>AACAAUGUUCAGUAAGUCAAAGACGAU | IDT | N/A |
| hs.Ri.DDB1.13.1<br>Top strand (5′→3′):<br>GCACAACCUACUUAUCAUUGACCAA<br>Bottom strand (5′→3′):<br>UUGGUCAAUGAUAAGUAGGUUGUGCAC | IDT | N/A |
| hs.Ri.DDB1.13.2<br>Top strand (5′→3′):<br>GACUUGAUUGAGAGUUUCCUGGATA<br>Bottom strand (5′→3′):<br>UAUCCAGGAAACUCUCAAUCAAGUCAC | IDT | N/A |
| hs.Ri.TRIM68.13.1<br>Top strand (5′→3′):<br>CCUUGCUACAGCAUUGGAACCAACA<br>Bottom strand (5′→3′):<br>UGUUGGUUCCAAUGCUGUAGCAAGGAC | IDT | N/A |
| hs.Ri.TRIM68.13.2<br>Top strand (5′→3′):<br>GAUGAAGUAGAUACAGUCUUCAGGA<br>Bottom strand (5′→3′):<br>UCCUGAAGACUGUAUCUACUUCAUCAU | IDT | N/A |
| hs.Ri.TRIM3.13.1<br>Top strand (5′→3′):<br>AUCAUUGUGGUCGACAACAAGUCTT<br>Bottom strand (5′→3′):<br>AAGACUUGUUGUCGACCACAAUGAUAU | IDT | N/A |
| hs.Ri.TRIM3.13.2<br>Top strand (5′→3′):<br>AGAAUGGCACAUAUGAGCUAGUGTA<br>Bottom strand (5′→3′):<br>UACACUAGCUCAUAUGUGCCAUUCUUG | IDT | N/A |
| hs.Ri.TRIM59.13.1<br>Top strand (5′→3′):<br>GCUUAUUCUGUACAUUGCUAAACAA | IDT | N/A |

| | | |
|---|---|---|
| Bottom strand (5′→3′): UUGUUUAGCAAUGUACAGAAUAAGCCC | | |
| hs.Ri.TRIM59.13.2 Top strand (5′→3′): CUCCAACUGGCAUUGAAUCUUUACC Bottom strand (5′→3′): GGUAAAGAUUCAAUGCCAGUUGGAGCA | IDT | N/A |
| **mRNA** | | |
| GFP mRNA | This Study | N/A |
| BZLF1 (ZTA) mRNA | This Study | N/A |
| **Recombinant DNA** | This Study | N/A |
| pCW57.1 | This Study | N/A |
| pCW57.1_DIPTAR | This Study | N/A |
| pCW57.1_DIPTAR[S297A] | This Study | N/A |
| pLEX307_DIPTAR | This Study | N/A |
| pLEX307_DIPTAR_2XFlag | This Study | N/A |
| pLEX307_DIPTAR[S297A]_2XFlag | This Study | N/A |
| pCW57.1_DIPTAR_U6_sgRNA | This Study | N/A |
| pCW57.1_HIF1α(FL)-CARD8 | This Study | N/A |
| pCW57.1_HIF1α(CT)-CARD8 | This Study | N/A |
| pCW57.1_HIF1α(ODD)-CARD8 | This Study | N/A |
| pCW57.1_HIF1α(CODD)-CARD8 | This Study | N/A |
| pCW57.1_HIF1α(ODD[P402A/P564A])-CARD8 | This Study | N/A |
| pCW57.1_HIF1α(CTAD)-CARD8 | This Study | N/A |
| pCW57.1_HIF1α(eGFP)-CARD8 | This Study | N/A |
| pCW57.1_HIF1α(ODD)-CARD8[S297A] | This Study | N/A |
| pLEX307_2XFLAG_HIF1α | This Study | N/A |
| pLEX307_HIF1α(ODD)-CARD8_2XFlag | This Study | N/A |
| pLEX307_HIF1α(ODD[P402A/P564A])-CARD8_2XFlag | This Study | N/A |
| pLEX307_CARD8_Flag | This Study | N/A |
| pLEX307_HIF1α(FL)-CARD8[S297A]_2XFlag | This Study | N/A |
| pLEX307_HIF1α(CT)-CARD8[S297A]_2XFlag | This Study | N/A |
| pLEX307_HIF1α(ODD)-CARD8[S297A]_2XFlag | This Study | N/A |
| pLEX307_HIF1α(CODD)-CARD8[S297A]_2XFlag | This Study | N/A |
| pLEX307_HIF1α(ODD[P402A/P564A])-CARD8[S297A]_2XFlag | This Study | N/A |
| pLEX307_eGFP-CARD8[S297A]_2XFlag | This Study | N/A |
| pLEX307_HIF1α(CTAD)-CARD8[S297A]_2XFlag | This Study | N/A |
| pLEX307_eGFP_Flag | This Study | N/A |
| pLEX307_DN-CUL2 | This Study | N/A |
| pCW57.1_ HIF1α(ODD)-CARD8_U6_sgRNA | This Study | N/A |
| pLenti-5HRE-GFP PuroR | Addgene | N/A |
| pCW57.1_BRD4-CARD8 | This Study | N/A |
| pLEX307_Ikβα-CARD8 | This Study | N/A |
| pLEX307_P53-CARD8 | This Study | N/A |
| pLEX304_MDM2 | This Study | N/A |
| pLEX304_MDM2(C464A) | This Study | N/A |

| | | |
|---|---|---|
| pgLAP1_GFP_CBU1314 | Shin | N/A |
| SuperPiggyBac Transposase | Shalem | N/A |
| PiggyBac CAS9-NLS-FLAG-P2A-mcherry | This Study | N/A |
| Bison Library | Busino | N/A |
| **Software and algorithms** | | |
| GraphPad Prism Version 11.02 | GraphPad Software | GraphPad Prism (RRID:SCR_002798) |
| Empiria Studio | Li-Cor Inc. | Empiria Studio (RRID:SCR_022512) |
| Cutadapt Version 5.2 | | (RRID:SCR_011841) |
| MAGeCK Version 0.5.9.4 | | (RRID:SCR_025016) |
